# Host innate immune response profiling reveals hidden viral infections across diverse animal species

**DOI:** 10.64898/2026.03.16.711951

**Authors:** Luca Nishimura, Hiroaki Unno, Kyoko Kurihara, Mai Suganami, Junna Kawasaki, Spyros Lytras, Kaho Okumura, Edward C Holmes, Jumpei Ito, Kei Sato

## Abstract

Virus discovery using RNA-seq data from wildlife and livestock offers a powerful strategy for identifying unknown pathogens with pandemic potential. However, conventional approaches rely on homology-based searches that have limited sensitivity for highly divergent viruses, are computationally intensive at scale, and cannot distinguish true infections from contamination. Viral infection induces interferon-stimulated genes (ISGs), key components of the frontline antiviral defense, and their expression serves as a robust indicator of viral infection. Here, we developed a host-response–based virus discovery framework that rapidly quantifies ISG expression and predicts viral infection status. Applying this framework to ∼210,000 RNA-seq data sets from diverse mammalian and avian species, we identified hidden viral infections across diverse hosts, including those caused by highly divergent viruses missed by a conventional approach. Our framework complements existing virus discovery strategies by adding host innate immune response context and enabling computationally efficient prescreening for scalable viral surveillance.

**Highlights:**

- Host response–based virus discovery in wildlife and livestock RNA-seq data
- Quantify interferon-stimulated gene (ISG) expression and predict viral infection
- Analysis of ∼210,000 RNA-seq data sets reveals hidden viral infections
- Detects highly divergent viruses and scalable viral surveillance through rapid prescreening

## Introduction

Most human viral infectious diseases arise through cross-species transmission from other animals, and such zoonotic spillover events continue to pose a major threat to global health^1^. The identification of viruses in wildlife and livestock reservoirs is therefore central to understanding spillover risk and viral evolution. The accumulation of large amounts of RNA-seq data in public repositories such as the Sequence Read Archive (SRA) has fueled a recent surge in virus discovery^2–6^. However, the volume of RNA-seq data generated each year is growing exponentially^7^. For example, approximately 170,000 mammalian and avian RNA-seq data sets had been deposited in the SRA by 2023, while approximately 40,000 data sets were released in 2024 alone. Because conventional virome analyses relying on *de novo* assembly and homology-based searches are computationally intensive, virus discovery strategies with lower computational costs are becoming a research imperative. In addition, homology-based searches—particularly those optimized for speed—are inherently limited in detecting infections caused by highly divergent viruses or those with few viral sequences. Nor can they determine whether any viral sequences detected represent genuine infections or arise from contamination^8^.

To overcome these challenges we developed a host innate immunity-centric strategy for detecting viral infection and identifying diverse viral pathogens. Viral infection activates interferon (IFN) pathways and induces IFN-stimulated genes (ISGs), many of which function as antiviral effectors^9^. Because this response is triggered by diverse viruses, ISG expression profiles can serve as a general indicator of viral infection, as shown in human transcriptomic studies^10–12^. As IFN responses are evolutionarily conserved across vertebrates^13,14^, this framework may be applicable to virus discovery across diverse animal taxa. Based on these principles, we developed the ISG Profiler and ISG-VIP to rapidly quantify ISG expression and predict viral infection status from mammalian and avian RNA-seq data (**Fig. 1A**). By applying this framework to approximately 210,000 publicly available RNA-seq data sets we identified previously hidden viral infections and uncovered novel, likely pathogenic viruses, providing new insights into viral evolution and future spillover risk.

**Fig. 1.**
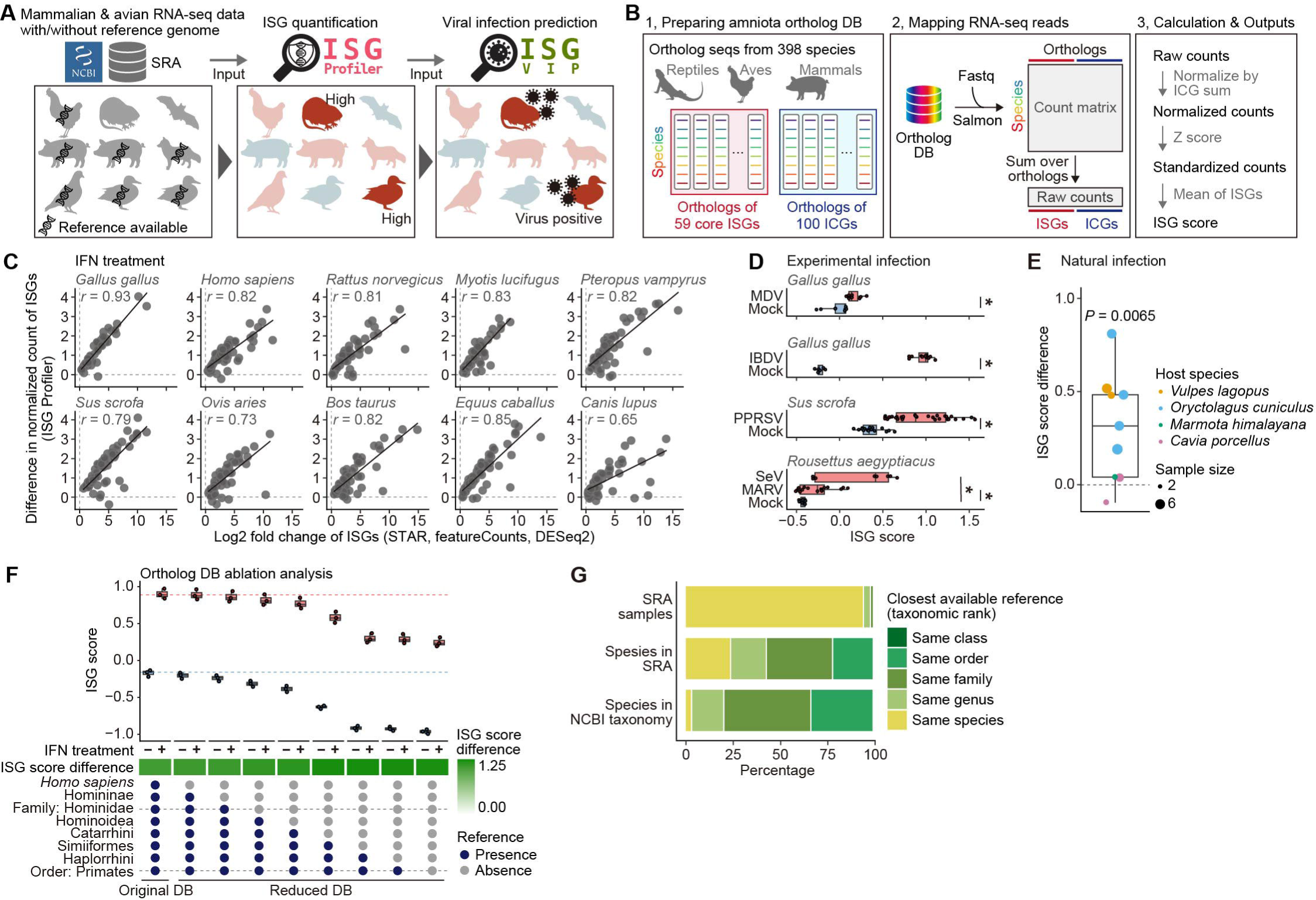
Development of ISG Profiler. A) Host IFN response–based virus discovery flamework applicable for diverse mammalian avian species. B) Overview of ISG Profiler. C) Induction levels of ISGs in response to type I IFN stimulation, estimated using ISG Profiler and a conventional RNA-seq pipeline. RNA-seq data from Shaw *et al.*^13^ were analyzed, in which cultured cells derived from diverse animal species were stimulated with type I IFN. Each dot represents a core ISG. Linear regression lines and Pearson correlation coefficients are shown. Sample information is listed in **Table S3**. D) Increase in ISG scores following experimental viral infection. Dots represent biological replicates, and boxplots show quartiles and Tukey’s fences. Asterisks indicate *P* < 0.05 (two-sided Welch’s t-test). MDV, Marek’s disease virus (MDV); IBDV, infectious bursal disease virus; PRRSV, porcine reproductive and respiratory syndrome virus; and SeV, Sendai virus. E) Changes in ISG scores associated with natural viral infection. RNA-seq data from lung samples of diverse animal species in which viral infections were detected^20,21^. The y axis shows differences in mean ISG scores between virus-infected and non-infected samples. Dots represent unique combinations of animal species and viral families. *P* values were calculated using two-sided one-sample *t*-tests. F) Ablation analysis to assess the robustness of ISG Profiler to limited genome availability. Top: ISG scores for IFN-treated and untreated samples calculated using the original and reduced databases; dashed lines indicate medians obtained with the original DB. Middle: Differences in mean ISG scores between IFN-positive and IFN-negative samples. Bottom: Composition of the ortholog DB variants, showing the taxonomic levels of reference sequences included in each DB. G) Availability of reference sequences across taxonomic levels. Proportions are shown for RNA-seq data sets registered in the SRA, unique animal species included in SRA and the NCBI taxonomy data set (https://www.ncbi.nlm.nih.gov/taxonomy).

## Results

### Overview of ISG Profiler

Although conventional RNA-seq analyses require species-specific reference sequences, such references are not always available when analyzing samples from diverse animal species. To overcome this challenge, we developed ISG Profiler, a framework that rapidly quantifies ISG expression at the ortholog level from mammalian and avian RNA-seq data without requiring species-specific reference sequences. The ISG Profiler first maps RNA-seq reads to a reference database comprising ortholog sequences derived from 398 amniote species for two gene sets: (i) 59 core ISGs, a set of experimentally validated conserved ISGs across 10 mammalian and avian species^13^, and (ii) 100 internal control genes (ICGs) (**Fig. 1B**). Mapped reads are then aggregated at the ortholog level to obtain raw counts. These counts are normalized by the total ICGs raw counts, log-transformed, and subsequently standardized using precomputed means and standard deviations for orthologs. Finally, an ISG score, representing the overall ISG expression state of each sample, is calculated as the mean of the standardized counts across all core ISGs. In summary, ISG Profiler outputs raw, normalized, and standardized counts for core ISGs and ICGs, along with an ISG score for each sample. On average, ISG Profiler processes standard paired-end 150-bp RNA-seq data in 3.9 minutes on a system equipped with eight 2.4 GHz CPU cores and 16 GB of RAM (**Fig. S1A**).

### Validation of expression quantification accuracy by ISG Profiler

To validate ISG Profiler, we first conducted benchmark analyses using an RNA-seq data set of IFN-treated cultured cells across ten mammalian and avian species^13^. To assess quantification of individual ISG induction, we compared differences in standardized counts between IFN-treated and untreated samples with log₂ fold-change values estimated by a conventional differential expression analysis pipeline^15–17^ (**Fig. 1C**). Strong correlations were observed across all species examined, supporting the accuracy of ISG Profiler. Consistent with this, IFN stimulation resulted in a substantial increase in ISG scores (**Fig. S1B**). In addition, the analysis of RNA-seq data from experimental viral infections confirmed that ISG Profiler detected marked elevations in ISG scores in infected samples (**Fig. 1D**)^18,19^. To further assess whether ISG Profiler captures ISG induction associated with natural viral infections, we analyzed RNA-seq data sets from lung tissues of wildlife and livestock in which viral infections had been identified in previous studies^20,21^; this revealed that ISG scores were generally higher in virus-positive samples than in virus-negative samples (**Figs. 1E and S1C**). Collectively, these results demonstrate that ISG Profiler robustly captures ISG induction associated with IFN stimulation, experimental viral infection, and natural viral infection.

### Quantification of ISG expression in species lacking reference genome information

To evaluate whether ISG Profiler can be applied to RNA-seq data from animal species lacking species-specific reference sequences, we conducted an ablation analysis in which portions of the ortholog reference database were systematically removed (**Fig. 1F**). Using RNA-seq data from IFN-treated human cells^13^, we showed that when reference sequences from species within the same family were available, ISG scores deviated only minimally from those obtained using the full reference database. A similar trend was observed for chicken (*Gallus gallus*) RNA-seq data (**Fig. S1D**). We next analyzed RNA-seq data from experimental viral infections in animal species for which reference sequences are available only at the genus or family levels; pronounced ISG upregulation was observed in all cases (**Fig. S1E**). These results suggest that reference sequences from phylogenetically related species at the family level are sufficient for unbiased ISG quantification by ISG Profiler. In addition, although species-specific references are not available for most mammalian and avian species recorded in the NCBI Taxonomy database, a comprehensive catalogue of known species, family-level references are available for 62.6% of these species (**Fig. 1G**). These results suggest that ISG Profiler is broadly applicable across diverse mammalian and avian taxa.

### ISG expression profiling of the 170k mammalian and avian RNA-seq samples

Next, we applied ISG Profiler to 168,438 mammalian and avian RNA-seq data sets registered in the NCBI SRA (hereafter referred to as the “170k RNA-seq” data set) (**Fig. 2A**). Human and mouse samples were excluded. Because we aimed to compare ISG expression profiles between virus-infected and noninfected samples, we also excluded data lacking assembled contig information in the Logan database, a large-scale resource providing *de novo* assemblies of public RNA-seq data sets^22^, which we subsequently used for viral detection. The resulting data set spans 997 species, 735 genera, 279 families, and 60 orders and comprises data from 6,234 BioProjects (**Figs. 2B and 2C**). In addition to tissue-derived samples, this data set includes RNA-seq data generated from swabs, fecal samples, and cultured cells.

**Fig. 2.**
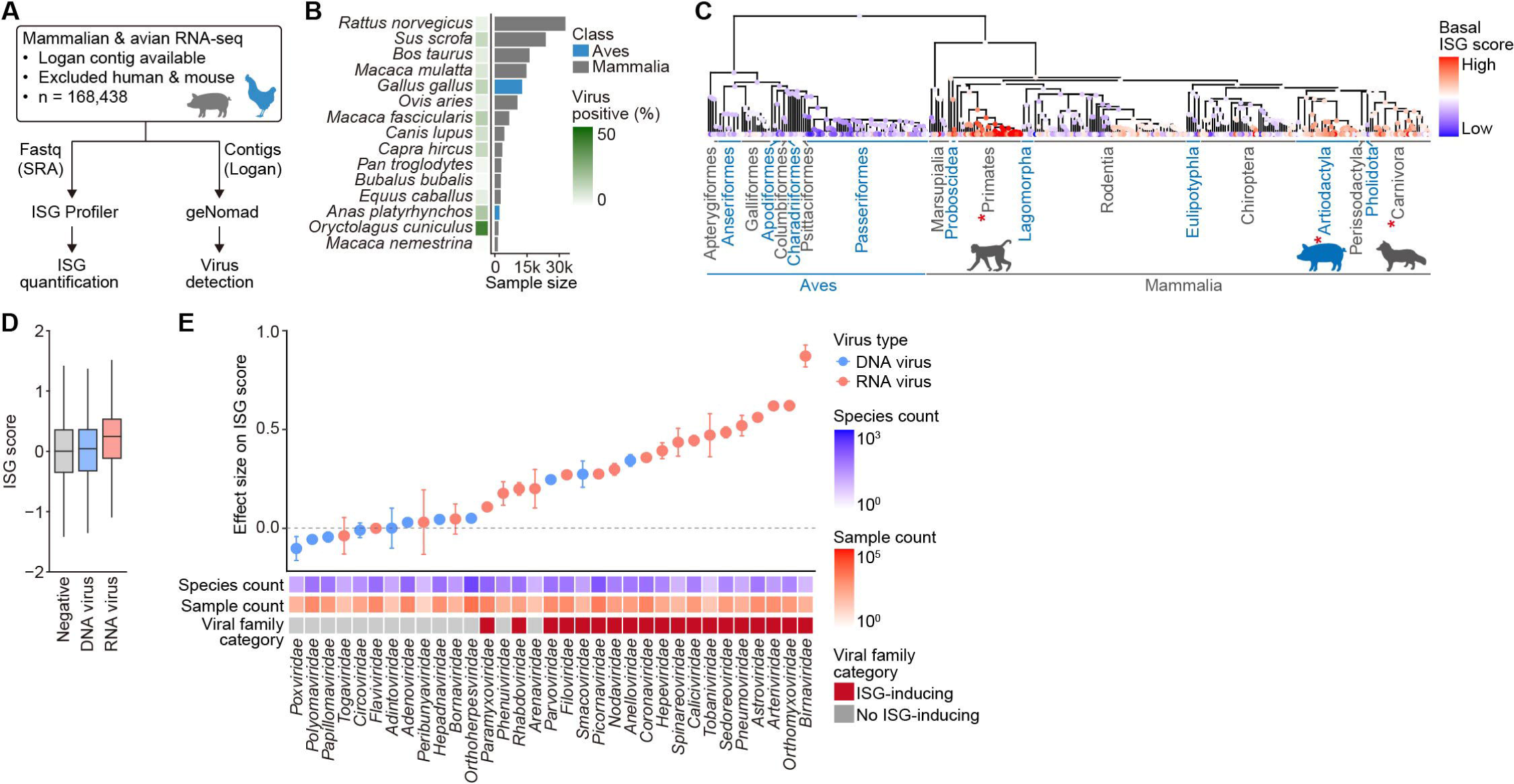
ISG profiling at the pan-SRA scale across host species and viral taxa. A) Overview of the 170k RNA-seq data set. B) Number of samples per animal species (bar) and virus-positive rates (heatmap). The top 15 species by sample count are shown. C) Basal ISG expression across animal species. The basal ISG score was defined as the mean ISG score in non-infected samples for each species. Species with more than five samples are shown. Inferred ancestral states of basal ISG scores are displayed at internal nodes of the species tree. Asterisks indicate significant elevation of basal ISG scores in the corresponding animal orders compared with other lineages (*P* < 0.05; two-sided Wald test in a linear mixed-effects model). D) Distributions of ISG scores in virus-negative samples and samples positive for DNA or RNA viruses. Boxplots show quartiles and Tukey’s fences. Samples co-infected with DNA and RNA viruses were excluded. E) Estimated effects of infection by each viral family on ISG scores. Estimates were obtained using a linear model adjusting for host species effects; points indicate estimated effects and error bars denote 95% confidence intervals. Bottom heatmaps show, from top to bottom, the number of host species and samples in which each viral family was detected, and classification of viral families as ISG-inducing. ISG-inducing viral families were defined as *P* < 0.01 or estimated effect size > 0.1.

In parallel, we assessed viral positivity in each RNA-seq data set by applying geNomad^23^, a widely used state-of-the-art virus detection tool that primarily relies on homology-based searches, to contigs reconstructed by Logan. This analysis detected virus-like contigs spanning diverse viral families in 10.5% of RNA-seq data sets (**Figs. S2A and S2B**). A substantial proportion of virus-positive samples originated from experimental infection studies, such as SARS-CoV-2 (*Betacoronavirus pandemicum*) infections in golden hamsters (*Mesocricetus auratus*) (**Figs. S2C and S2D**). Furthermore, some virus-like contigs detected by geNomad—particularly those assigned to *Orthoherpesviridae*—showed top hits to non-viral sequences in BLASTx searches, suggesting that a fraction of these detections may represent false positives (**Figs. S2A and S2E**). Although these caveats should be considered, we nevertheless constructed the 170k data set comprising ISG expression profiles and viral positivity information.

### Comparison of ISG expression profile across animal hosts and viral taxa

Differences in basal ISG expression levels are known to be associated with variation in viral susceptibility and disease outcomes across animal species^24^. We therefore examined ISG scores in uninfected samples, referred to as basal ISG scores, across host species. Basal ISG scores were significantly higher in several placental mammalian orders, including Primates, Artiodactyla, and Carnivora, compared to other lineages (**Figs. 2C and S2F**). A similar pattern was observed for the expression of STAT1, a central regulator of ISG transcription^14^, supporting elevated basal ISG expression in these lineages. Consistent with their polyphyletic distribution, ancestral state reconstruction indicated that elevated basal ISG expression evolved convergently across these mammalian orders (**Fig. 2C**).

We next compared ISG expression levels across viral taxa. Samples positive for RNA viruses exhibited higher ISG scores overall, whereas samples positive for DNA viruses showed ISG scores comparable to those of virus-negative samples (**Figs. 2D and S2G**). To quantify differences in ISG responses across viral families while adjusting for host species effects, we constructed a linear model to estimate the effect of infection by each viral family on ISG scores (**Fig. 2E**). Infection by RNA virus families generally exerted positive effects on ISG scores, whereas most DNA virus families did not. Notable exceptions among RNA viruses included the *Bornaviridae*, *Togaviridae*, and *Flaviviridae* which did not show significant positive effects on ISG scores. Bornaviruses are known to establish persistent infections that are often associated with limited innate immune activation, whereas togaviruses and flaviviruses encode viral proteins that potently antagonize IFN signaling pathways^25–27^. In contrast, among DNA viruses, several single-stranded DNA virus families, including the *Parvoviridae*, *Anelloviridae*, and *Smacoviridae*, showed positive effects on ISG scores. These patterns remained consistent when ISG scores were compared between infected and uninfected samples within each host species, supporting the robustness of this pattern (**Fig. S2H**). Hereafter, viral families that exhibited positive effects on ISG scores in this statistical modeling analysis are defined as ISG-inducing viral families (annotated in **Fig. 2E**).

### Development and validation of ISG-VIP

Next, we developed a machine learning model, termed the ISG-based viral infection predictor (ISG-VIP), to predict the viral infection status of individual RNA-seq samples based on ISG expression profiles quantified by ISG Profiler together with host taxonomic information (**Fig. 3A, left**). ISG-VIP incorporates expression levels of 100 ICGs in addition to 59 ISGs, as ICG expression patterns may help the model infer and adjust for baseline ISG expression levels influenced by factors such as tissue differences. Because not all viral families induce robust ISG expression (**Fig. 2E**), ISG-VIP was trained using geNomad-based viral detections from ISG-inducing viral families. We trained and evaluated ISG-VIP using the 170k RNA-seq data set, applying a five-fold cross-validation scheme (**Fig. 3A, right**). ISG-VIP achieved higher predictive performance than simple logistic regression models based on the ISG score alone or on the expression of individual ISGs combined with host taxonomic information (**Fig. 3B**). Predictive performance was particularly high for viral families and host species with high virus-positive rates (**Fig. S3A**).

**Fig. 3.**
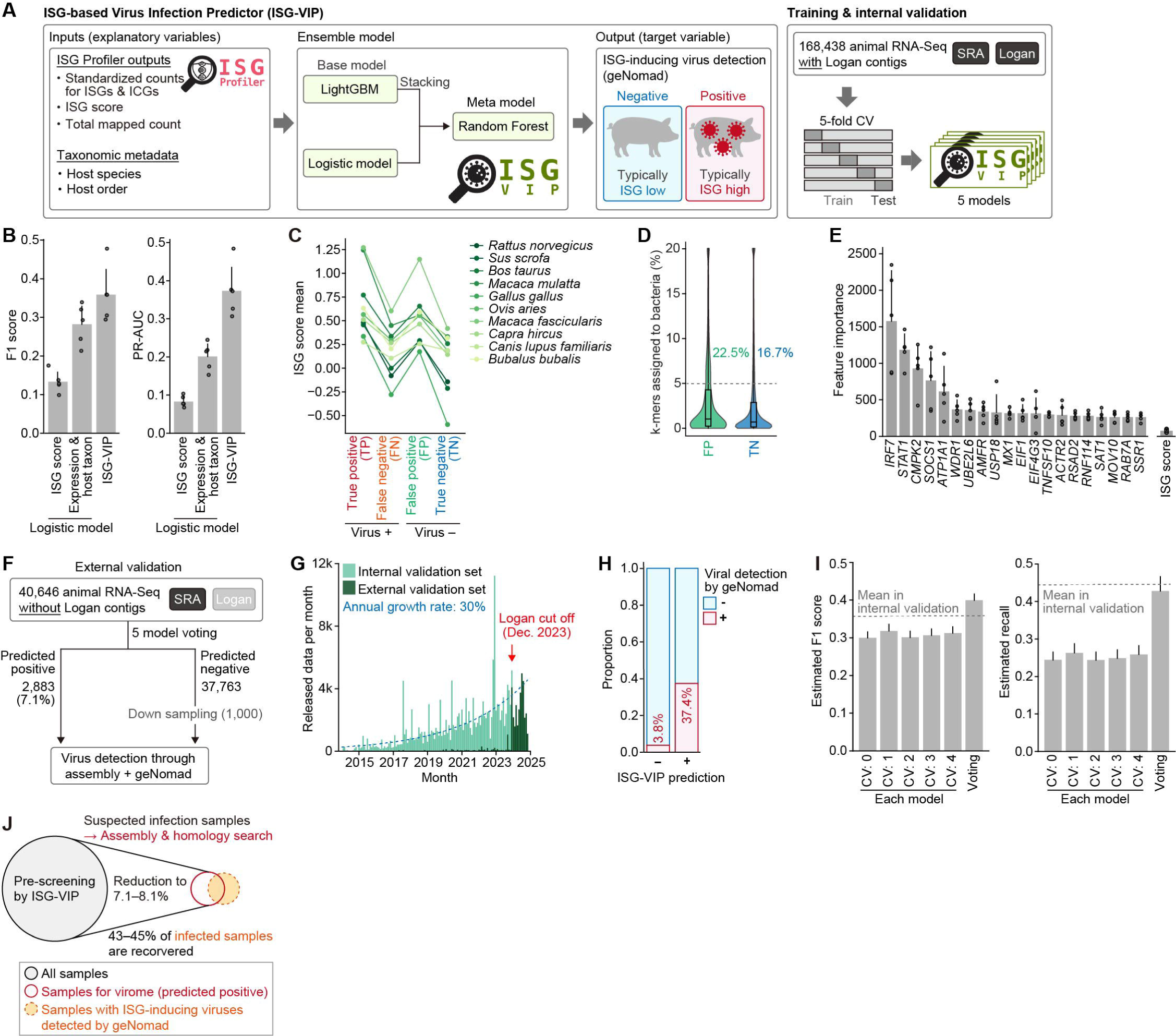
Development of the viral infection prediction model ISG-VIP. A) Architecture of the ISG-VIP model (left) and training scheme (right). B) F1 score (left) and PR-AUC (right) of ISG-VIP. Values for individual folds (dots), mean values (bars), and standard deviations (error bars) are shown. Results from logistic regression models are shown for comparison. C) Mean ISG scores in TP, FN, FP, and TN fractions of the confusion matrix. Results are shown for the 10 host species with the largest numbers of virus-positive samples. D) Proportion of bacterial-like sequences (k-mers) in each RNA-seq data measured by STAT^51^. Boxplots show quartiles and Tukey’s fences. The proportions of samples exceeding 5% bacterial k-mers (dashed line) are indicated. E) Feature importance of individual variables in the LightGBM component of ISG-VIP. F) Scheme for external validation of ISG-VIP prediction accuracy. G) Distribution of RNA-seq release dates in the internal and external validation data sets. A stacked bar plot shows the number of samples released each month. An exponential trend line fitted to the monthly counts using Poisson regression is overlaid. H) Comparison of detection frequencies of ISG-inducing viral families between predicted-positive and predicted-negative fractions in the external validation. I) Estimated F1 score and recall for individual models and for majority voting (bars), with standard errors. Performance values are estimates, as predicted-negative samples were down-sampled prior to virus detection. J) Efficient virus discovery workflow based on ISG expression profiling. The expected proportion of samples subjected to virome analysis, together with the expected recovery rate of virus-infected samples, is shown based on the positive prediction rate and recall obtained in internal and external validations.

To better understand the strengths and limitations of ISG-VIP, we characterized samples in each category of the confusion matrix: true positives (TP), false positives (FP), true negatives (TN), and false negatives (FN). Among virus-positive samples, ISG scores were higher in the TP group than in the FN group, whereas among virus-negative samples, ISG scores were higher in the FP group than in the TN group (**Fig. 3C**). These results suggest that ISG-VIP preferentially detects viral infections associated with strong ISG induction, while occasionally misclassifying non-infected samples with elevated ISG expression as virus-positive. Indeed, a subset of FP samples exhibited higher proportions of bacterial sequences than TN samples, suggesting that some FP predictions likely reflect bacterial infections accompanied by ISG induction (**Fig. 3D**).

To identify key marker genes contributing to viral infection prediction, we analyzed feature importance and SHAP (SHapley Additive exPlanations) values for input variables in ISG-VIP (**Figs. 3E and S3B**). IRF7 and STAT1 were the strongest individual contributors to infection prediction among all ISGs and surpassed the predictive performance of the overall ISG score, supporting the importance of considering individual gene expression levels in addition to the ISG score.

To further assess the robustness of ISG-VIP prediction, we performed external validation using mammalian and avian 40,646 RNA-seq data sets in the SRA for which contig data were not available in the Logan database (hereafter referred to as the “41k RNA-seq” data set) (**Fig. 3F**). Because the Logan project analyzed RNA-seq data released only up to December 2023, most data in this external validation set were released in 2024 (**Fig. 3G**)^22^. We applied models obtained from five-fold cross-validation to the 41k RNA-seq data set and predicted infected samples by majority voting. We then reconstructed contigs for these samples using a pipeline consistent with Logan^23,28,29^ and assessed viral infection status using geNomad. As a control, we applied the same analysis to 1,000 samples downsampled from the predicted-negative group.

In the external validation, ISG-VIP predicted 2,883 samples (7.1% of the total) as infection-positive, of which viruses were detected in 37.3% (**Fig. 3H**). In contrast, viruses were detected in only 3.4% of predicted infection-negative samples. The expected F1 score was estimated at 0.40, exceeding those observed in internal validation (**Fig. 3I**). The recall, representing the proportion of virus-infected samples detected, was estimated at 0.43, supporting the robustness of ISG-VIP prediction.

Taken together, these results suggest that ISG-VIP prescreening facilitates a computationally efficient virome workflow. In this workflow, all RNA-seq data are first screened with ISG-VIP, followed by computationally intensive assembly and homology searches only for samples predicted to be infected (**Fig. 3J**). Based on the predicted positive rate and recall from external and internal validation (**Figs. 3F, 3I, and S3C**), this workflow reduces the number of samples subjected to virome analysis to 7.1–8.5% while recovering 43–45% of virus-infected samples, demonstrating its practical effectiveness (**Fig. 3J**).

### Identification of novel viruses using ISG-VIP

To further demonstrate the utility of the ISG profiling–based workflow for virus discovery (**Fig. 3J**), we attempted to identify novel viruses in the 41k RNA-seq data set. Nearly all of these data were released in 2024 and were not included in previous large-scale virome studies that comprehensively analyzed animal RNA-seq data in the SRA (**Fig. 3G**)^2–6^.

For virus discovery, we utilized a BLASTx-based pipeline. In this pipeline, we first performed BLASTx searches against the NCBI non-redundant (nr) protein database and retained contigs whose top BLASTx hits corresponded to eukaryotic viral sequences. To further exclude potential non-viral sequences, additional BLASTn filtering against the NCBI core nucleotide (nt) database was applied. Although this pipeline is slower than geNomad-based virus detection, it provides higher sensitivity and specificity. Accordingly, contigs detected by this pipeline are referred to as virus-derived (rather than virus-like) contigs.

Identified virus-derived contigs were then grouped into infection events, defined as unique combinations of samples and viral genera, yielding 2,441 infection events (**Fig. 4A**). To prioritize high-confidence infection events involving viruses with low similarity to known viruses, we selected events with a cumulative hit length >200 amino acids and a mean top-hit sequence identity < 95%, hereafter referred to as candidate novel virus infection events.

**Fig. 4.**
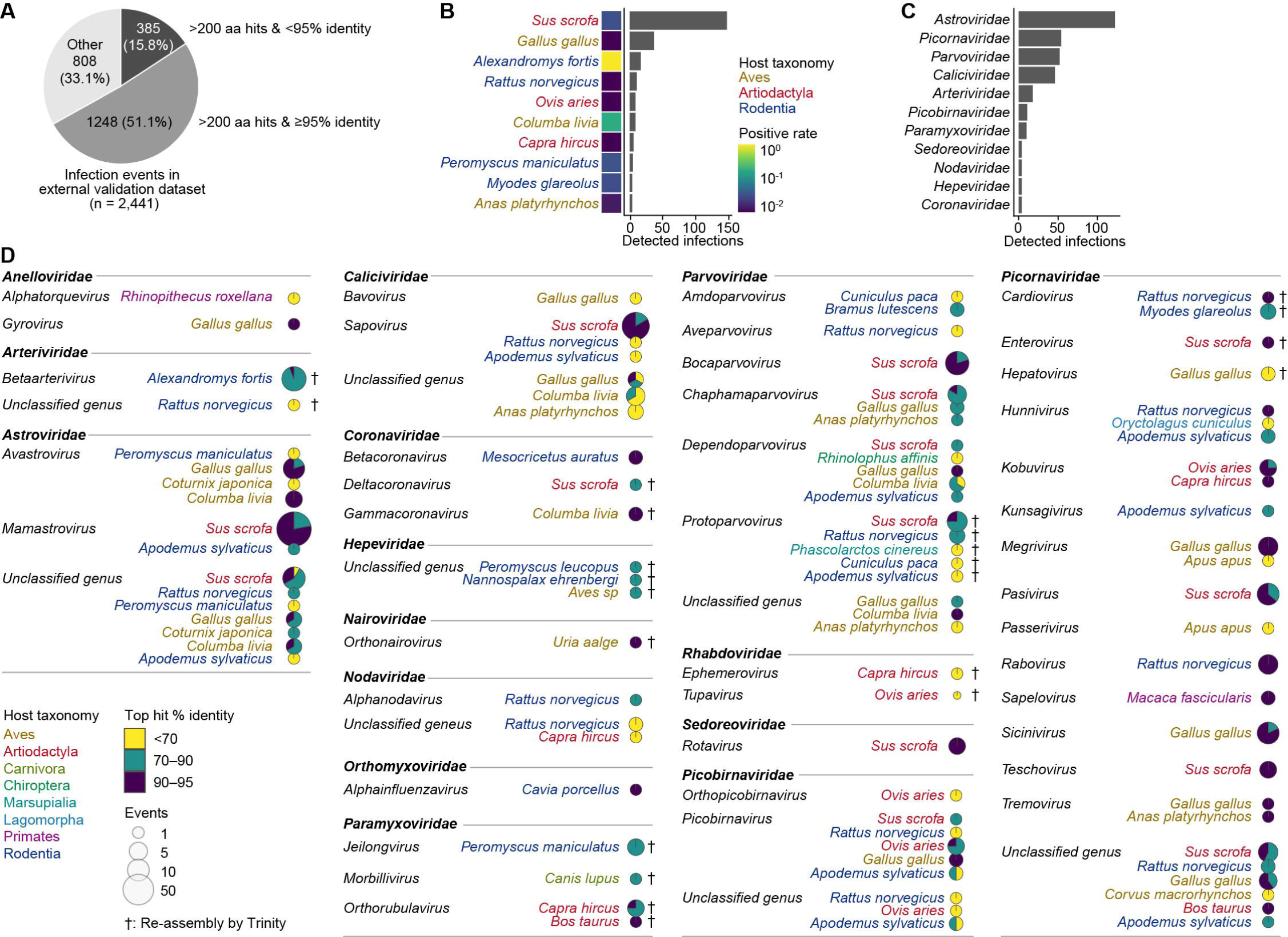
Discovery of novel viruses from the external validation data set. A) Composition of viral infection events detected by BLASTx against the NCBI non-redundant (nr) protein sequence database in the external validation (40k) data set. Detected viral infection events are listed in **Table S4**. B) Number of viral infection events by host species. The positive rates of candidate novel viral infections are shown as a heatmap. C) Number of viral infection events by viral family. D) Number of candidate novel virus infection events detected in each host species. Pie size indicates the number of events, and segment proportions indicate BLASTx top-hit sequence identity categories. Viral families known to infect only fungi, plants, or invertebrates were excluded. Infection events subjected to detailed inspection after high-quality Trinity^30^ reassembly are indicated by daggers.

In total, 385 candidate novel virus infection events were identified across diverse host species, including livestock, laboratory model animals, rodents, and birds (**Fig. 4B**). Many infection events were assigned to the *Astroviridae*, *Picornaviridae*, *Parvoviridae*, *Caliciviridae*, and *Arteriviridae* (**Fig. 4C**). These events comprised 97 unique combinations of viral genera and host species (**Fig. 4D**). Because the contig sequences generated by the Logan pipeline were often highly fragmented, we reassembled contigs using Trinity^30^, which is slower but capable of reconstructing higher-quality contigs (**Figs. S3D and S3E**). This reassembly was performed for RNA-seq data sets with infection events assigned to viral genera that include known human, zoonotic, or livestock pathogens, to enable further validation (daggers in **Fig. 4D**). The reconstructed high-quality contigs were then mapped to reference viral genomes, confirming that they covered substantial portions of viral genomes and suggesting that these sequences were genuinely virus-derived (**Fig. S4**). Together, these results demonstrate that the ISG profiling–based workflow enables the identification of a broad range of viruses across diverse host species.

### Detection of viral infections missed by a conventional method using ISG-VIP

Although geNomad is a widely used state-of-the-art virus detection tool^23^ and was used for ground-truth labeling in ISG-VIP training, its predictions are not always accurate. Consequently, the FP fraction may include true viral infections missed by geNomad (**Fig. 5A**). To test whether ISG-VIP can identify such missed cases, we searched for viruses in the FP fraction of one internal cross-validation data set (CV: 0) using the BLASTx-based method described above. All FP samples (n = 2,169) were analyzed, with TP (n = 536), FN (n = 769), and 1,000 randomly selected TN samples (from n = 22,142) used as controls. Viruses were detected by BLASTx in 85% of TP samples; most of the remaining 15% were excluded because their top BLASTn hits corresponded to non-viral sequences (**Figs. 5B and S5A**). Importantly, viruses were detected in 16.8% of FP samples but in only 6.4% of TN samples, indicating that ISG-VIP captures a substantial fraction of infected samples missed by geNomad. Comparison of the mean BLASTx top-hit sequence identity per sample showed that highly divergent matches (≤70% identity) accounted for 13.7% of FP samples, compared with 6.8% in the TP fraction, suggesting enrichment of novel viruses in the FP fraction (**Fig. 5C**). In addition, the cumulative BLASTx hit length per sample was substantially shorter in the FP fraction than in the TP fraction (**Fig. 5D**). Consistent with this, among the virus-derived contigs in this analysis, approximately half of the contigs ≤1,000 bp could not be identified by geNomad (**Fig. S5B**). These results highlight that ISG-VIP combined with BLASTx searches can detect viral infections associated with highly divergent or fragmented virus-derived contigs that are often missed by geNomad.

**Fig. 5.**
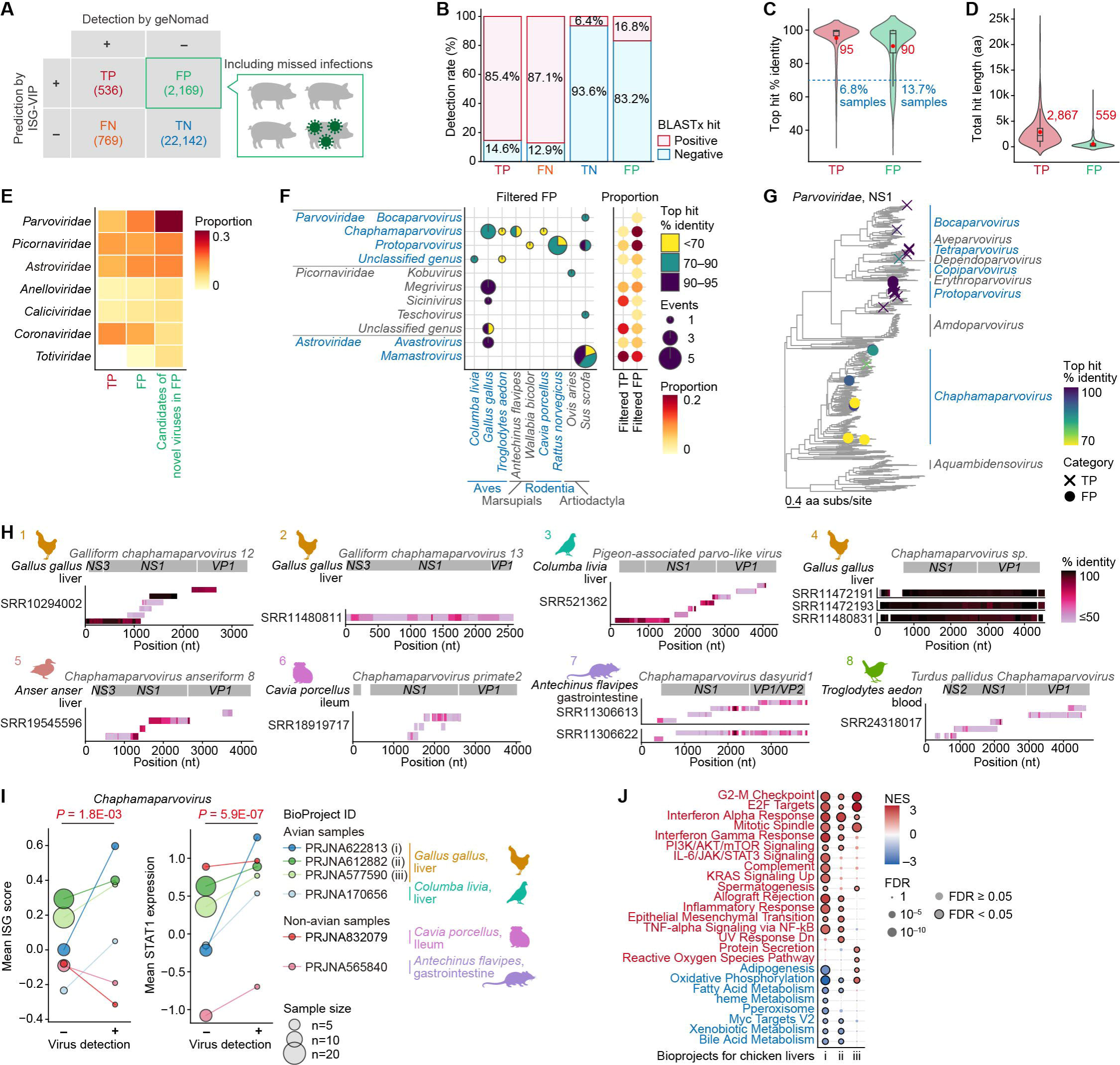
Detection of previously overlooked viral infections. A) Confusion matrix and overlooked viral infections. The FP fraction includes both genuine false positives and viral infections missed by geNomad. Results for the Cross validation (CV):0 data set are shown. B) Viral detection rates by BLASTx in each fraction. ISG non-inducing viral families were not counted as positive. C) Distribution of BLASTx mean top-hit sequence identity. When multiple infection events were detected within a sample, the per-sample mean was used. Boxplots show quartiles and Tukey’s fences, with red dots and numbers indicating means. In addition, the proportions of samples with ≤70% identity are shown in blue. D) Cumulative length of BLASTx hit regions per sample. E) Proportions of viral families among detected infection events in each fraction. The plot includes viral families detected at a frequency ≥5% among candidate novel virus infection events in the FP fraction. Detected viral infection events are listed in **Table S4**. F) Numbers of candidate novel virus infection events detected in each host species. Results for the FP fraction are shown for viral genera within the *Parvoviridae*, *Picornaviridae*, and *Astroviridae*. The heatmap on the right shows the proportions of viral genera among candidate novel virus infection events in the FP and TP fractions. G) Maximum likelihood tree of *Parvoviridae* NS1 proteins. In addition to NS1 sequences identified from the TP and FP fractions, all parvoviral NS1 protein sequences from the NCBI Virus database were included after quality control and clustering at 90% sequence identity. BLASTx top-hit sequence identity and data source (TP or FP) of the identified sequences are annotated. H) Mapping of *Chaphamaparvovirus* contigs to the reference genome by TBLASTx. High-quality contigs obtained by Trinity reassembly were mapped to the near-complete reference genome that were most closely related to the BLASTx top-hit virus. Rows represent contigs, and color indicates animo acid sequence identity. Gene annotations of the reference genome are shown. The numbers in the upper left of each plot correspond to the clusters defined in the phylogenetic analysis shown in **Fig. S5F**. Details of reassembled high-quality contigs are listed in **Table S5**. I) ISG induction associated with *Chaphamaparvovirus* infection. Mean ISG score (left) and standardized STAT1 counts (right) are shown for virus-positive and virus-negative groups. Virus-negative samples were selected from the same BioProjects under comparable conditions. Only viruses for which comparable virus-negative samples were available were included in the analysis. Dot size indicates sample size. *P* values were calculated using mixed-effects models adjusting for BioProject effects. Sample information is listed in **Table S3**. J) Gene expression changes associated with *Chaphamaparvovirus* infection in chicken liver samples. Two-group comparisons were performed using DESeq2 for each BioProject, followed by GSEA^59^ based on Wald statistics using MSigDB gene sets^60^. The top 10 positively and negatively enriched terms based on the normalized enrichment score (NES) are shown for terms with FDR < 0.05 in at least one BioProject, colored red and blue, respectively. BioProject: PRJNA622813 (i), PRJNA612882 (ii), PRJNA577590 (iii).

The ground-truth labels used for ISG-VIP training do not account for viral families that do not induce significant ISG expression, here defined as ISG non-inducing viral families. Consequently, when such viruses infect samples in the absence of coinfecting viruses, these cases appear in the FP fraction if detected by ISG-VIP and in the TN fraction otherwise. To examine whether ISG-VIP can detect infections caused by these viral families, we compared their detection frequencies between the FP and TN fractions (**Fig. S5C**). Significant enrichment in the FP fraction was observed for the majority of ISG non-inducing viral families, most prominently *Orthoherpesviridae* and *Hepadnaviridae*. These findings suggest that infections by ISG non-inducing viral families can nonetheless be detected by ISG-VIP when individual infections trigger ISG responses.

To identify the frequently missed viruses by geNomad we extracted candidate infection events involving novel viruses (>200 amino acid cumulative hit length and <95% mean identity in BLASTx) from the FP fraction (**Fig. 5E**). Consistent with the external validation results, many infection events involving the *Parvoviridae*, *Picornaviridae*, and *Astroviridae* were detected in the FP fraction. Notably, the *Parvoviridae*—a single-stranded DNA virus family characterized by short genomes and lacking a polymerase gene—showed the strongest enrichment in the FP fraction. Within the *Parvoviridae*, the genera *Chaphamaparvovirus* and *Protoparvovirus* were particularly enriched in this fraction (**Fig. 5F**). Although *Chaphamaparvovirus* is a relatively recently characterized parvoviral genus, officially defined by ICTV in 2020^31^, our phylogenetic analyses showed that it is the most diverse genus within the *Parvoviridae* (**Figs. 5G and S5D**). In addition, the *Chaphamaparvovirus* sequences identified here showed markedly lower similarity to known viruses than sequences from other genera (**Fig. 5G**), suggesting that the genetic diversity of *Chaphamaparvovirus* remains largely underrepresented in current virus databases. We also confirmed that geNomad failed to identify any of *Chaphamaparvovirus*-derived contigs identified in this study, even when using the latest versions of geNomad (v1.11.2) and its database (v1.9) (**Fig. S5E**).

### Novel viruses causing hepatitis-like transcriptome signatures in chickens

We next performed a detailed biological characterization of viruses newly identified by our workflow. Using ISG-VIP with BLASTx searches, we identified 11 *Chaphamaparvovirus* infection events supported by NS1 hits from chickens, wild birds, marsupials, and rodents (**Fig. 5H**). Phylogenetic analysis revealed that the identified sequences clustered into eight distinct groups representing diverse evolutionary lineages (**Fig. S5F**). For these viruses, particularly those identified in birds, ISG scores and STAT1 expression levels were higher in virus-detected samples than in non-detected samples derived from the same BioProjects under comparable conditions, suggesting that these represent genuine infections (**Fig. 5I**).

Notably, most of these chaphamaparvoviruses were detected in samples derived from liver tissues of chickens or other avian species (**Fig. 5H**). Previous studies have also reported associations between *Chaphamaparvovirus* infection and histopathologically diagnosed hepatitis in chickens and pheasants^32,33^. To access the potential association between hepatitis and the diverse identified *Chaphamaparvovirus* lineages in chickens, we performed transcriptome-wide differential expression analysis on infected chicken liver samples (**Figs. 5J and S5G**). Gene set enrichment analysis (GSEA) showed upregulation of pathways related to antiviral responses (Interferon Alpha Response), inflammation (IL-6/JAK/STAT3 Signaling and Inflammatory Response), cytotoxic T cell infiltration (Allograft Rejection), and liver remodeling (Epithelial–Mesenchymal Transition). In contrast, pathways associated with hepatic metabolic functions, including Xenobiotic Metabolism and Bile Acid Metabolism, were downregulated, suggesting liver dysfunction. Together, these transcriptional signatures observed in infected chicken liver samples are consistent with virus-associated hepatic inflammation.

We identified another example of a potential hepatitis-associated virus in chickens. In the external validation (41k) data set the genus *Hepatovirus*, which comprises mammalian viruses including human hepatitis A virus, was detected in liver samples from chickens in France and China (**Fig. 6A**). Although known avian hepatitis viruses phylogenetically related to mammalian *Hepatovirus* (e.g., duck hepatitis virus type 1) exist, they are classified as a distinct genus, *Avihepatovirus* (**Fig. 6B, left**). Nevertheless, recent metagenomic studies have reported *Hepatovirus*-like sequences in fecal samples from aquatic birds^34–37^. Our phylogenetic analysis showed that these avian *Hepatovirus*-like sequences form a distinct clade separate from *Hepatovirus* and exhibit genetic diversity comparable to that of related genera (**Figs. 6B and S6A**). In particular, we first identified the *Hepatovirus* sequences from chicken liver samples clustered within this novel avian clade (**Fig. 6B, right**). This result suggests spillover from aquatic birds into chickens, followed by spread within chicken populations and hepatic infection. GSEA showed robust upregulation of antiviral and inflammatory pathways in one infected sample, whereas another infected sample exhibited broad downregulation of genes associated with liver function, possibly reflecting distinct disease stages (**Figs. 6C and S6B**). Although based on limited evidence, these transcriptional signatures are consistent with virus-associated hepatic inflammation.

**Fig. 6.**
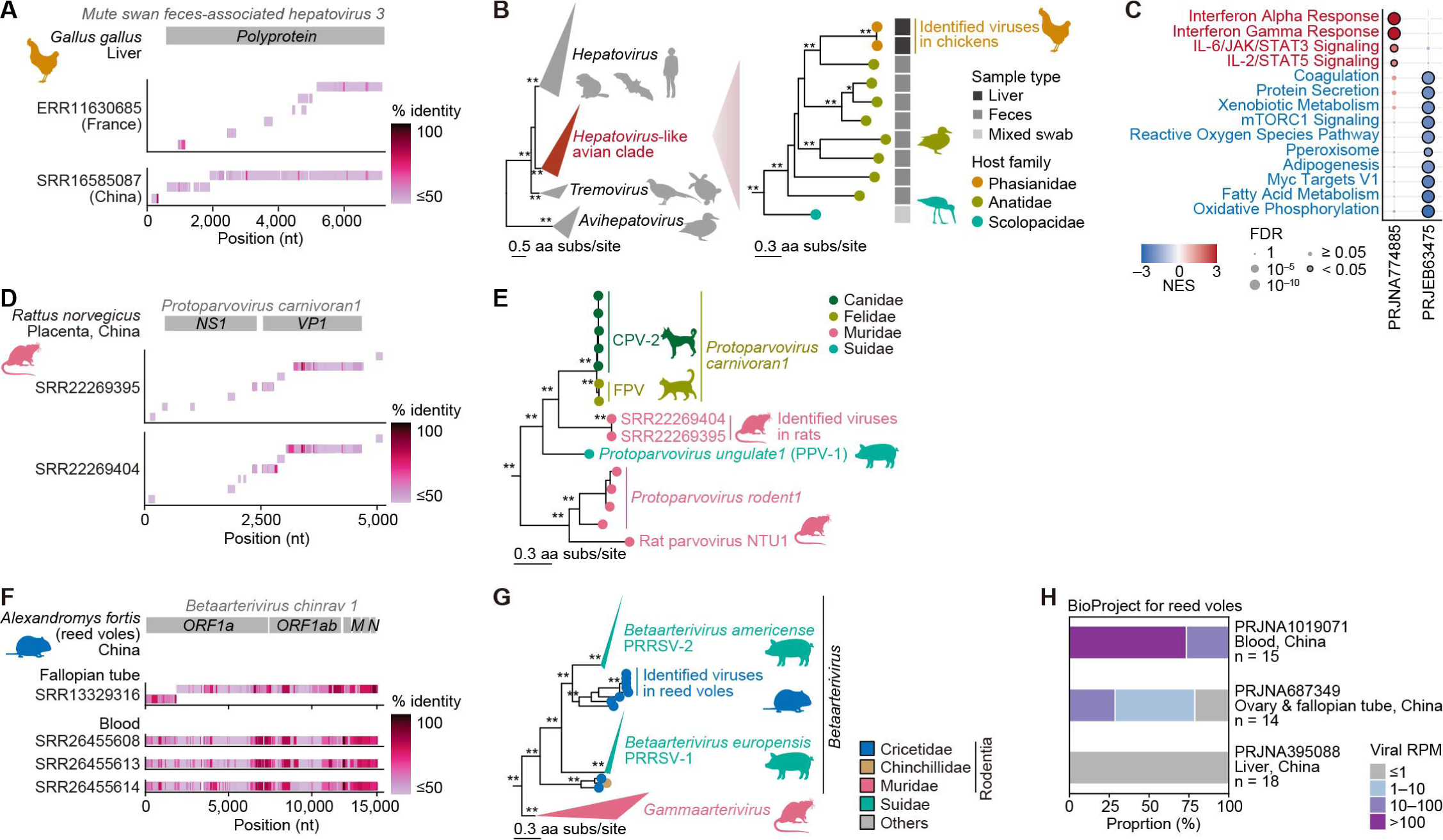
Characterization of newly identified viruses. A) Mapping of *Hepatovirus* contigs identified from chicken liver samples by TBLASTx. B) Maximum likelihood phylogenetic tree based on polyprotein sequences of the genus *Hepatovirus* and related picornavirus genera. The tree was midpoint rooted. The right panel shows a subtree of the avian *Hepatovirus*-like clade, including the viruses identified in this study. C) Gene expression changes associated with *Hepatovirus* infection in chicken liver samples, assessed by GSEA. Sample information is listed in **Table S3**. D) Mapping of *Protoparvovirus* contigs identified from rat placenta samples by TBLASTx. E) Maximum likelihood tree of the identified viruses and related viruses, including feline panleukopenia virus [FPV] and canine parvovirus type 2 [CPV-2]) and *Protoparvovirus ungulate1*, also known as porcine parvovirus 1 (PPV-1). The full phylogenetic tree of *Protoparvovirus* VP1 sequences was estimated using IQ-TREE 3^56^, from which the presented subtree was extracted. Double and single asterisks indicate ultrafast bootstrap support values of >95 and >90, respectively. F) Mapping of *Betaarterivirus* contigs identified from reed vole samples by TBLASTx. G) Maximum likelihood tree of *Betaarterivirus* with *Gammaarterivirus* used as outgroups. The full phylogenetic tree of *Arteriviridae* ORF1 sequences was estimated, from which the presented subtree was extracted. H) Detection frequency of the novel *Betaarterivirus* in RNA-seq data sets derived from reed voles. All SRA BioProjects with at least two samples were analyzed, and samples with >1 reads per million (RPM) were considered virus-positive.

### Evolutionary origin of highly pathogenic carnivore and porcine protoparvoviruses

We identified a novel *Protoparvovirus* missed by geNomad in the 170k data set (**Fig. 5F**). This virus was detected in rat placental samples derived from a single BioProject (**Fig. 6D**). The genus *Protoparvovirus* includes several highly pathogenic viruses, such as porcine parvovirus 1 (PPV-1; *Protoparvovirus ungulate1*) and *Protoparvovirus carnivoran1*, which includes feline panleukopenia virus (FPV) and canine parvovirus type 2 (CPV-2). PPV-1 forms a clade with FPV and CPV-2. Our phylogenetic analysis showed that the *Protoparvovirus* sequences identified in rats (Muridae) fell within this clade rather than clustering with the Muridae-associated viruses that were the outgroup to this clade (**Fig. 6E**). This phylogenetic topology, together with ancestral host inference (**Fig. S6C**), suggests a Muridae origin for the clade of highly pathogenic protoparvoviruses including PPV-1, FPV, and CPV-2.

### High prevalence of PRRSV-related *Betaarterivirus* in reed voles

In the 41k data set a particularly high frequency of virus detection was observed in reed voles (*Alexandromys fortis*), largely attributable to the identification of novel *Betaarterivirus* (**Figs. 4B and 4D**). This virus is related to porcine reproductive and respiratory syndrome virus (PRRSV) and was detected in two BioProjects from China (**Fig. 6F**). PRRSV was first recognized in the late 1980s and is now one of the most economically important pathogens in swine production^38^. PRRSV comprises two species, PRRSV-1 (*Betaarterivirus europensis*) and PRRSV-2 (*Betaarterivirus americense*), which are thought to have arisen through two independent spillover events from rodent viruses into pigs^38,39^. Phylogenetic analysis showed that the reed vole viruses identified form a single clade within the sister clade of PRRSV-2, which comprises viruses derived from Cricetidae hosts (the rodent family that includes reed voles) (**Fig. 6G**). Notably, three BioProjects containing ≥2 reed vole samples are available in the SRA database, two of which—originating from Hunan Province, China—exhibited viral positivity rates exceeding 75% (**Fig. 6H**). These results suggest that a *Betaarterivirus* lineage closely related to viruses responsible for two independent global swine epidemics circulates in a reed vole population in China, likely at high prevalence.

## Discussion

Although infection prediction based on the expression of host genes primarily including ISGs has been developed in humans^10–12^, no general framework applicable across diverse animal species has been established. Here, we filled this gap by developing a scalable host IFN-response–based virus discovery framework comprising ISG Profiler and ISG-VIP (**Fig. 1 and 3**). The framework processes RNA-seq data within an average of 4 minutes and is applicable across diverse mammalian and avian species, including those lacking species-specific reference sequences. This framework can serve as a powerful complement to existing virome discovery pipelines because it can identify viruses missed by conventional homology–based methods and provide additional information on host innate immune responses to detected viruses. Indeed, application of this framework enabled the identification of many novel viruses (**Fig. 4**), including a chicken *Chaphamaparvovirus* and *Hepatovirus* associated with hepatitis-like transcriptomic signatures (**Fig. 5**), a rat *Protoparvovirus* providing insights into the evolutionary origins of known animal pathogens (**Fig. 6**), and *Betaarterivirus* detected in a reed vole colony potentially posing elevated spillover risk (**Fig. 6**). These results highlight the utility of the framework.

In virome analysis, the possibility that any viral sequences detected represent contamination must always be considered. In addition to the consistency between the sampled species and known host ranges, elevated ISG expression—a canonical host response to viral infection—provides supporting evidence for a *bona fide* infection. Accordingly, viral sequences recovered from samples classified as infection-positive by ISG-VIP, which infers robust ISG induction (**Fig. 3C**), are more likely to reflect true infections. Thus, ISG Profiler and ISG-VIP can provide supporting evidence that the sequences detected represent genuine infections rather than contamination and are of viral origin. In this way, our approach adds a new layer of information—host innate immune responses—to conventional virome analyses, enabling a more nuanced interpretation of viral infections.

In addition, we propose a computationally efficient virome workflow in which RNA-seq data are first prescreened using ISG Profiler and ISG-VIP, followed by detailed virome analysis including contig assembly and homology-based searches (**Fig. 3J**). This workflow reduces the number of samples subjected to computationally intensive virome analyses to 7.1–8.5% while recovering 43–45% of virus-infected samples. Although this strategy may miss viral infections that do not induce ISG responses, it enables sustained monitoring of the rapidly expanding number of newly generated data sets, which is increasing at an annual rate of approximately 30% (**Fig. 3G**).

This study has several limitations. First, ISG-VIP is conceptually unable to detect viruses that do not induce ISG expression. Some pathogenic viruses, such as members of the *Togaviridae* and *Flaviviridae,* possess strong IFN-antagonistic activity and therefore elicit minimal ISG responses, limiting their detectability by this approach (**Fig. 2E**). Nevertheless, infections caused by ISG non-inducing viral families—which were not considered during ISG-VIP training—were detectable when individual infections triggered ISG induction, suggesting that this framework is applicable beyond predefined ISG-inducing viral families (**Fig. S5C**). Second, ISG induction is not specific to viral infections and can also be triggered by other pathogens, including bacteria, which may lead to false-positive predictions (**Fig. 3D**). However, this observation also suggests that ISG-VIP could potentially be repurposed for detecting non-viral pathogens that induce IFN responses. Finally, the lack of appropriate virus-negative control samples in many public RNA-seq data sets, together with uncertainty in virus detection, may bias estimates of infection-associated expression changes.

Despite these limitations, our study demonstrates that host gene expression–based virus discovery provides a powerful framework for viral surveillance in wildlife and livestock. This framework can be readily extended beyond ISGs to include other gene sets, such as inflammatory responses, apoptosis, or cellular stress pathways, potentially enabling detection of viruses that do not strongly induce ISGs. Moreover, integrating host expression patterns with virus discovery may allow simultaneous inference of viral pathogenicity and immune evasion strategies, advancing toward a unified platform for infection detection and functional characterization. Because ISG Profiler can in principle be applied to samples containing RNA from multiple host species, such as environmental RNA, monitoring ISG expression in wastewater or farm environments may enable early detection of outbreaks caused by unknown viruses. Continued development of host-response–based virus discovery approaches thus holds promise for establishing next-generation viral surveillance systems that integrate early detection with risk assessment of emerging pathogens.

## Methods

### Data download

FASTQ files for RNA-seq data sets in the NCBI SRA were downloaded using the SRA Toolkit (v3.1.1) with the prefetch and fastq-dump commands (https://github.com/ncbi/sra-tools/wiki/01.-Downloading-SRA-Toolkit). FASTQ files publicly available from the National Genomics Data Center (NGDC) were downloaded using wget (v1.19.5). A complete list of SRA accessions analyzed in this study is provided in **Data sets 1–4**. Reference genome sequences (FASTA) and gene annotations (GTF) for host animals were obtained from the NCBI FTP site (ftp.ncbi.nlm.nih.gov/genomes/all/GCF/). The reference genomes and annotations used in this study are summarized in **Table S1**. Taxonomic information for host animals and viruses was obtained from the NCBI taxonomy dump (release date: June 5, 2025; ftp://ftp.ncbi.nlm.nih.gov/pub/taxonomy/taxdump.tar.gz).

Orthology relationships of host genes were extracted from the NCBI FTP (ftp.ncbi.nlm.nih.gov/gene/DATA/gene_orthologs.gz; release date: June 10, 2024) and were linked to the corresponding RefSeq identifiers using gene2refseq.gz (release date: June 12, 2024). mRNA isoform sequences registered in NCBI RefSeq were retrieved using isoform accession numbers and the Entrez efetch utility from the Biopython package (v1.83). A list of human genes (including protein-coding and noncoding genes) was obtained from the HGNC BioMart database (https://biomart.genenames.org; downloaded on September 24, 2024). To quantify gene expression variability across human tissues, we downloaded the GTEx Analysis V8 bulk tissue expression matrix^40^ (https://storage.googleapis.com/adult-gtex/bulk-gex/v8/rna-seq/GTEx_Analysis_2017-06-05_v8_RNASeQCv1.1.9_gene_reads.gct.gz).

A host species phylogeny (**Fig. 2C**) was obtained from TimeTree (downloaded on September 10, 2025; https://timetree.org)^41^. Because phylogenetic information for some of the 398 amniote species included in the ortholog database is not available in TimeTree, an alternative species tree covering all included species was generated using the ETE Toolkit v4 (retrieved on December 19, 2024)^42^. This tree depicts topology only and does not contain branch length information (**Fig. S1G**)

To obtain RNA-seq data sets from mammals and birds, we mined the NCBI SRA (excluding human and mouse) on November 14, 2024 using the following search terms, used in a previous study^3^: [(“Mammalia”[Organism] OR “Mammals”[All Fields]) AND (“biomol rna”[Properties] AND “library layout paired”[Properties] AND “filetype fastq”[Properties]) NOT (“Homo sapiens”[Organism]) NOT (“Mus musculus”[Organism])] or [(“Aves”[Organism] OR “Aves”[All Fields]) AND (“biomol rna”[Properties] AND “library layout paired”[Properties] AND “filetype fastq”[Properties])]. FASTQ files were downloaded for SRA accessions returned by these searches. We then manually curated the downloaded accessions and removed data sets that were erroneously included and did not originate from mammals or birds.

Pre-assembled contig data sets provided by Logan^22^ (v1.1) were downloaded as FASTA files using the AWS CLI with the following parameters: aws s3 cp s3://logan-pub/c/<accession>/<accession>.contigs.fa.zst. --no-sign-request. Amino acid sequences and associated metadata of known viruses used for phylogenetic analysis were obtained from NCBI Virus (https://www.ncbi.nlm.nih.gov/labs/virus/vssi/#/).

### Mapping RNA-seq reads to host reference genomes

Adapters and low-quality bases were removed from the downloaded RNA-seq reads using fastp (v1.0.1) with default parameters^43^. Processed reads were mapped to the corresponding host reference genome using STAR (v2.7.11a) ^15^. STAR was run with the following parameters:

--outFilterMultimapScoreRange 1 --outFilterMultimapNmax 10 --outFilterMismatchNmax 5 - -alignIntronMax 500000 --alignMatesGapMax 1000000 --sjdbScore 2 --alignSJDBoverhangMin 1 --genomeLoad NoSharedMemory --limitBAMsortRAM 0 --outFilterMatchNminOverLread 0.33 --outFilterScoreMinOverLread 0.33 --sjdbOverhang 100 --outSAMstrandField intronMotif --outSAMattributes NH HI NM MD AS XS --outSAMunmapped Within --outSAMtype BAM Unsorted --outSAMheaderHD @HD VN:1.4. Gene-level read counts were quantified from BAM files using featureCounts (v2.0.6)^16^.

### Differential expression analysis

Differential expression between two conditions was assessed using DESeq2 (v1.44.0 or v1.50.2)^17^ based on the gene count matrix generated from the reference-genome mapping pipeline described above. For each gene, log_2_ fold change, Wald statistics, and false discovery rates (FDRs) adjusted by the Benjamini–Hochberg method were computed.

### Mapping RNA-seq reads to the mRNA isoform sequence database

For mapping RNA-seq reads to the mRNA isoform sequence database, we evaluated Salmon (v1.10.1; --validateMappings)^44^, KMA (v1.4.15; -nc -na -nf -ef)^45^, Kallisto (v0.50.1; -l 100 -s 30)^46^, STAR (same parameters as above), and Bowtie2 (v2.5.2; default parameters)^47^. For STAR- and Bowtie2-based mappings, BAM files were sorted using Samtools (v1.19)^48^.

Read counts were extracted from tool-specific output files as follows: NumReads from Salmon (quant.sf), est_counts from Kallisto (abundance.tsv), mapped read counts from KMA (*.mapstat), and exon-level gene counts from STAR and Bowtie2 using featureCounts (v2.0.6; -t exon -g gene_id -fracOverlap 0.25).

### Overview of ISG Profiler

We developed ISG Profiler, a computational tool that rapidly quantifies ISG expression at the ortholog level without requiring species-specific reference genomes for each analyzed host. ISG Profiler first maps RNA-seq reads to a reference sequence database constructed from ISGs and ICGs derived from 398 amniote species (**Fig. 1B**). Mapped reads are then aggregated at the ortholog level to generate raw counts, which are normalized by the total raw counts of ICGs and subsequently log-transformed to obtain normalized expression values for each ortholog. Next, standardized counts are computed using the mean and standard deviation of ortholog expression levels estimated from 168,438 RNA-seq data registered in the SRA. Finally, an ISG score representing the overall ISG expression state of each sample is calculated as the mean of the standardized counts across all ISGs. In summary, ISG Profiler takes processed FASTQ files as input and outputs raw, normalized, and standardized counts for ISGs and ICGs, together with the ISG score for each sample.

As ISGs, we used 59 previously defined core ISGs^13^ that show high evolutionary conservation across mammals and birds. As ICGs, we selected 100 genes that are highly conserved and expressed among amniotes but exhibit minimal expression variability across animal species, across tissues, and in response to IFN stimulation. For read mapping, we adopted Salmon, which showed the highest correlation with expression estimates obtained using conventional approaches among the evaluated methods. Because the core ISG set includes genes conserved only in eutherian mammals or only in mammals, these ISGs are automatically excluded from quantification when analyzing marsupial or avian data sets (**Fig. S1F**). In addition to ortholog-level quantification integrated across animal species, ISG Profiler implements a species-resolved mode that estimates expression levels for each ortholog within individual host species (**Fig. S1G**). This mode is, in principle, applicable to samples with unknown or mixed host origin, such as fecal or environmental samples, by identifying the host species in which ISGs are upregulated.

### Selection of ISGs

We started with the 62 core ISGs defined by Shaw et al.^13^, which were experimentally validated as an evolutionarily conserved IFN-responsive gene set across cultured cells from 10 mammalian and avian species. We excluded *HLA* genes because orthologs were poorly conserved outside humans. We further filtered out genes showing outlier behavior in the IFN stimulation RNA-seq data set from Shaw et al. Specifically, for each gene, we regressed the replicate-averaged expression estimated by a conventional reference-genome-based pipeline (STAR + featureCounts; log10(RPM+1)) against the replicate-averaged standardized counts produced by ISG Profiler. Genes with an absolute Studentized residual ≥ 2 were considered outliers and excluded from the analysis, resulting in the removal of *RNF19B* (**Fig. S1H and Table S2**). In addition to *RNF19B* and *HLA* genes, *EIF2AK2* was excluded from the final list of core ISGs in ISG Profiler as it was an outlier. In an early prototype of ISG Profiler based on 30 internal control genes (ICGs) (**Fig. S1I**). However, in the final version of ISG Profiler using 100 ICGs, *EIF2AK2* was no longer detected as an outlier (**Fig. S1H**). Because ISG expression levels are strongly correlated with one another, the exclusion of a single ISG (i.e., *EIF2AK2*) is unlikely to substantially affect ISG profiling or downstream virus infection prediction.

### Selection of ICGs

We selected ICGs as genes that are highly conserved across species and highly expressed, yet exhibit minimal expression variability across host species, across tissues, and between IFN-stimulated and unstimulated conditions. ICG selection was performed as follows (**Fig. S1J**).

First, we identified genes conserved across the 10 species analyzed by Shaw et al.^13^ (*Gallus gallus*, *Homo sapiens*, *Rattus norvegicus*, *Myotis lucifugus*, *Pteropus vampyrus*, *Sus scrofa*, *Ovis aries*, *Bos taurus*, *Equus caballus*, and *Canis lupus*).

Specifically, among human genes, we retained genes for which one-to-one orthologous relationships were confirmed for all 10 species in NCBI gene_orthologs.gz.

Second, to select genes minimally responsive to IFN stimulation, we calculated the log2 fold change for each gene upon IFN stimulation in each species using RNA-seq data from Shaw et al., and then computed the absolute mean log2 fold change across species. Genes with an absolute mean log2 fold change across species ≤ 0.25 were retained.

Third, to identify genes that are highly expressed and stable across species, we calculated the mean expression level (log2(RPM+1)) and coefficient of variation (CV) across the 10 species, and rank-normalized these metrics (higher mean expression = higher rank; lower CV = higher rank). To further ensure stability across tissues and individuals, we performed the same procedure using GTEx Analysis V8 bulk tissue expression data and obtained rank-normalized mean expression and CV for each gene. Finally, we retained genes ranked within the top 10% across all four criteria (mean expression and CV in Shaw et al.; mean expression and CV in GTEx). For these genes, we computed the average rank-normalized score and selected the top 100 genes as ICGs. The selected ICGs are listed in **Table S2**.

### Construction of the ortholog database

The ortholog database was constructed by extracting orthologous mRNA isoform sequences corresponding to the 59 core ISGs and 100 ICGs from 398 amniote species. First, we extracted sequences orthologous to the human 59 core ISGs and 100 ICGs from 398 amniote entries in NCBI gene2refseq.gz. We then retrieved isoform accession numbers for these genes gene2refseq.gz and downloaded the corresponding sequences using Entrez efetch. The final database contains 151,856 mRNA isoform sequences (**Data set 9**).

### Selection of the read mapping tool

To select the mapping tool used in ISG Profiler, we benchmarked five mapping methods (Salmon, KMA, Kallisto, STAR, and Bowtie2) using RNA-seq data set from the IFN stimulation experiments across 10 species in Shaw et al. For each method, we computed standardized counts for each gene and calculated the difference in mean standardized counts between IFN-treated and untreated samples. We then computed the Spearman correlation between these values and the corresponding gene-wise log2 fold changes estimated by a conventional reference-genome-based pipeline (STAR + featureCounts + DESeq2) (**Fig. S1K**). Salmon showed the highest correlation and was therefore adopted as the mapping tool in ISG Profiler.

### ISG expression profiling of public RNA-seq data sets

We generated ISG expression profiles for mammalian and avian RNA-seq data sets in the NCBI SRA, excluding human and mouse. FASTQ files were processed using fastp with default parameters to remove adapter sequences and low-quality bases. Samples with a total raw ICG count ≤ 10,000 were excluded from downstream analyses. In total, 209,084 RNA-seq samples were analyzed. The list of data sets analyzed is provided in **Data sets 1–8**.

### Identification of virus-like contigs using geNomad

Among the RNA-seq data sets for which ISG expression profiles were obtained, 168,438 samples had pre-assembled contigs available from Logan v1.1^22^. To identify virus-like contigs, we extracted contigs ≥ 500 bp and annotated them using geNomad v1.8.0^23^ with the geNomad database v1.7 (options: --min-score 0.7 --cleanup) (database URL: https://zenodo.org/records/10594875). We filtered geNomad outputs using the following thresholds: virus_score ≥ 0.80, marker_enrichment ≥ 1.5, and n_hallmarks ≥ 1. We further excluded contigs assigned to bacteriophage-associated taxa (*Caudoviridae*, *Inoviridae*, *Lavidaviridae*, and *Microviridae*), *Retroviridae* (which frequently include endogenous retrovirus-derived hits), *Megaviricetes* and *Microviridae* (for which direct infection of vertebrates has not been reported), and unclassified sequences. Details of virus-like contigs identified in the 170k data set are summarized in **Data set 10**.

To validate the accuracy of our virome analysis pipeline (Logan contigs + geNomad), we compared our virus detection results with those reported by Kawasaki et al.^3^. This earlier work performed comprehensive RNA virus discovery using BLASTx on mammalian and avian RNA-seq data deposited in the SRA up to year 2019 (n = 46,359), which overlaps with our data set (**Fig. S2I**). Among samples classified as RNA virus–positive by Kawasaki et al., approximately 74% were also classified as RNA virus-positive in our analysis, whereas approximately 4% of samples classified as RNA virus-negative by Kawasaki et al. were classified as RNA virus-positive in our analysis. Collectively, these results indicate that virus detection using geNomad achieved sufficient sensitivity and specificity and is appropriate for downstream analyses.

### Validation of virus-like contigs detected by geNomad using BLASTx

For virus-like contigs detected by geNomad, we performed homology searches against NCBI non-redundant (nr) protein database (release date: May 5, 2025) using DIAMOND BLASTx (v2.1.8.162)^49^ and computed the percent identity to the top-hit protein. BLASTx analysis revealed that approximately 40% of the sequences classified as *Orthoherpesviridae* by geNomad had no top hits to known *Orthoherpesviridae* sequences, suggesting the presence of false positives (**Fig. S2A**). Among these sequences, approximately 80% showed top hits to host protein sequences, most frequently collagen alpha-3, Bcl-2, and polyadenylate-binding protein, indicating that geNomad likely misclassified these host-derived sequences as *Orthoherpesviridae* sequences (**Fig. S2E**).

The presence of numerous false-positive detections within *Orthoherpesviridae* may therefore lead to an underestimation of the effect of this viral family on ISG induction. Indeed, when these false-positive contigs were excluded and ISG scores were compared between *Orthoherpesviridae*-positive and -negative samples, a clear increase in ISG scores associated with *Orthoherpesviridae* infection was observed, whereas no such increase was apparent when infection status was defined solely based on geNomad classification (**Fig. S2J**). However, to maintain consistency in the main analyses presented in **Fig. 2**, we retained the geNomad classifications in subsequent analyses. Consequently, *Orthoherpesviridae* was not classified as an ISG-inducing viral family (**Fig. 2E**) and was not considered during ISG-VIP training. Nevertheless, ISG-VIP was still able to detect *Orthoherpesviridae* infection events (**Fig. S5C**), suggesting that this issue had limited impact on the overall results.

### Calculation of per-species basal ISG scores and ancestral state reconstruction

To compare basal ISG expression levels across host species, for each species with at least five virus-negative samples we calculated the median ISG score across virus-negative samples and defined this value as the per-species basal ISG score. We then scaled the per-species basal ISG scores and performed ancestral state reconstruction by treating the scaled basal ISG score as a continuous trait using the squared-change parsimony method implemented in the R package castor (v1.8.4)^50^.

### Estimating the effects of viral families on ISG scores and identifying ISG-inducing viral families

To compare ISG induction across viral families, we fitted a linear model in which the ISG score of each sample was the response variable and the detected viral family and host species were explanatory variables. The model was applied to 168,438 RNA-seq data sets. By analyzing regression coefficients for viral families, we estimated the effect size of infection by each viral family on ISG scores while adjusting for differences across host species. Statistical significance was assessed using Wald tests. Viral families with effect size > 0.1 and *P* < 0.01 were defined as ISG-inducing viral families.

### Development of ISG-VIP

ISG-VIP is a binary classification model that predicts, for each RNA-seq sample, the presence or absence of viral infection by ISG-inducing viral families detected by geNomad. As input features, we used standardized counts of ISGs and ICGs, the ISG score, the normalized total number of mapped reads, and one-hot encoded host taxonomic information (species and order). We included host order information in addition to species because, for some animal species, the number of samples was too small to reliably estimate species-specific effects. In contrast, ISG expression patterns tend to show consistent, order-level trends, so incorporating host order helps capture shared host biology and improves generalization (**Fig. S2F**). ISG-VIP is a stacking-based ensemble model. We used LightGBM and logistic regression as base learners and a random forest as the meta-learner. The model was implemented in Python (v3.12.7) using scikit-learn (v1.8.0) and lightgbm (v4.6.0). For training, we used 168,438 RNA-seq data sets for which contig information was available in the Logan database (the “170k data set”).

### Cross-validation design and preprocessing

We used 5-fold cross-validation for model training and performance evaluation. First, the full data set was split into five outer folds using scikit-learn’s StratifiedGroupKFold. To obtain a model that generalizes to unseen BioProjects, samples were grouped by BioProject ID so that samples from the same BioProject were not split across training and evaluation sets.

To prevent information leakage, within each outer fold the statistics required for feature standardization (mean and standard deviation) were estimated using only the training split, and these statistics were then used to transform both the training and evaluation splits. In addition, host categories (species and order) with fewer than five occurrences were treated as rare categories and merged into an “unknown” category during one-hot encoding within each outer fold.

The positive-class proportion in the training data used in this study was 5.4%, indicating class imbalance. Therefore, within each cross-validation split, we applied SMOTE from imblearn (v0.12.3) to the training data only to oversample the minority class. This was intended to mitigate learning bias and performance degradation caused by class imbalance.

### Hyperparameter optimization

Hyperparameter optimization was performed using Optuna (v4.2.1). For each outer fold, we further split the training data into five inner folds and conducted inner cross-validation during optimization. As the optimization objective, we used the sum of (i) the median F1 score computed across animal species and (ii) the PR-AUC computed over the entire training data set. For the per-species F1 score, we considered only species with at least five samples in the inner-training split and at least 200 samples in the full data set. We performed optimization for each base model as well as the stacking meta-model, with 50 trials for the base models and 50 trials for the meta-model.

For the LightGBM base model, we optimized learning_rate (0.01–0.1; log scale), num_leaves (50–100), min_data_in_leaf (10–40), feature_fraction (0.8–1.0), bagging_fraction (0.8–1.0), bagging_freq (1–3), lambda_l1 (1e-8–1.0; log scale), and lambda_l2 (1e-8–1.0; log scale).

For the logistic regression base model, we optimized C (0.01–10.0; log scale) and solver (“lbfgs”, “liblinear”).

For the random forest meta-model, we optimized n_estimators (350–600), max_depth (1–4), min_samples_split (25–90), min_samples_leaf (25–90), max_features (“sqrt”, “log2”, None), max_samples (0.7–0.9), ccp_alpha (0.002–0.02), and class_weight (“balanced”, “balanced_subsample”).

### Determination of the prediction threshold

The threshold for converting predicted scores into binary labels was determined separately for each outer fold by selecting the threshold that maximized the F1 score using predictions from the inner cross-validation. Candidate thresholds were selected from a grid that divided the range 0–1 into 100 bins. The F1 curve was smoothed using a moving average (window size = 5), and the threshold corresponding to the maximum of the smoothed curve was chosen. The selected threshold was then applied to the evaluation split of the corresponding outer fold to obtain the final classification results.

### Quantification of bacterial-derived k-mers

To estimate the degree of bacterial sequence contamination in each RNA-seq sample we analyzed publicly available NCBI STAT data that hierarchically assign k-mers in sequencing reads to taxonomic categories^51^. We retrieved the corresponding STAT results for the RNA-seq data sets analyzed in this study using NCBI SRA BigQuery (https://www.ncbi.nlm.nih.gov/sra/docs/sra-bigquery/) and extracted the proportion of k-mers assigned to the taxonomic category Bacteria.

### Analysis of feature importance

To interpret the contribution of each feature to infection prediction, we computed feature importance for the LightGBM base classifier using the lightgbm package. In addition, we computed SHAP values on the training data using shap (v0.48.0) and visualized the results.

### External validation of ISG-VIP

We performed external validation using 40,646 RNA-seq data sets for which contigs were not available in the Logan database. The five models obtained from 5-fold cross-validation were applied to each sample, and the final predicted label was determined by majority vote.

For samples predicted to be positive, contigs were re-assembled using the same pipeline as in Logan, and geNomad was used to determine the presence or absence of viral infection. As a control, we performed contig assembly and geNomad-based assessment for 1,000 samples randomly selected from the predicted-negative group. Details of virus-like contigs identified in the external validation data set are provided in **Data set 11**.

### Contig assembly using the Logan pipeline, MEGAHIT, or Trinity

For assembly using the Logan pipeline, processed FASTQ files generated after trimming with fastp (v1.0.1) were used as input. Unitigs were first constructed using Cuttlefish2 (v2.2.0)^28^ with a k-mer size of 31. Contigs were then generated from the extracted unitigs using Minia3 (v3.2.5)^45^ with the following options: -skip-bcalm -skip-bglue -redo-links. Contig assembly using MEGAHIT (v1.2.9)^52^ and Trinity (v2.15.1)^30^ was performed on the same fastp-processed FASTQ files with default parameters.

### Estimation of the F1 score in external validation

In the external validation, infection status was assessed by geNomad for all samples in the predicted-positive group, whereas in the predicted-negative group infection status was assessed only for a randomly selected subset (1,000 data). Therefore, TP and FP can be directly observed, but FN and TN for the full population cannot be directly observed. Consequently, the F1 score cannot be computed directly from these data. We therefore estimated the false-negative rate and true-negative rate based on the random sample drawn from the predicted-negative group. Assuming random sampling, the proportions of FN and TN observed in the sample were treated as unbiased estimators of the corresponding population proportions. We multiplied these estimated proportions by the size of the predicted-negative population to obtain the expected values of FN and TN in the population. Finally, we computed the expected F1 score using the observed TP and FP and the expected FN and TN.

Next, to quantify uncertainty in the estimated F1 score, we estimated a 95% confidence interval. Because sampling from the predicted-negative population was performed without replacement, we assumed that the number of false negatives in the sample follows a hypergeometric distribution. Because TP and FP were fully observed, they were treated as fixed (error-free) values, and the size of the predicted-negative population (FN + TN) was also treated as known and fixed. Thus, we assumed that uncertainty in the F1 score arises solely from the estimation error of FN.

The F1 score is defined as:

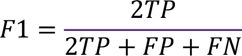

Using the delta method, we approximated the variance as:

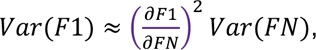

where Var(FN) was estimated from the hypergeometric distribution. Using the resulting standard error, we computed a 95% confidence interval based on a normal approximation.

### Identification of viral-derived contigs by BLASTx

To detect viral-derived contigs more sensitively than geNomad, we performed a comprehensive search against the NCBI RefSeq and nr protein databases using DIAMOND BLASTx (v2.1.12)^49^. To reduce computational cost, we adopted a two-stage search strategy.

First, we performed BLASTx against RefSeq viral proteins (download date: May 5, 2025) and extracted virus-like contigs satisfying e-value < 1e-4 and ORF length ≥ 50 amino acids. Next, to remove contigs that show greater similarity to non-viral proteins, we performed a second BLASTx search against the comprehensive NCBI nr protein database (download date: May 5, 2025) and extracted contigs satisfying the same criteria (e-value < 1e-4 and ORF length ≥ 50 amino acids).

For each contig, we obtained the top-hit subject based on bit score and assigned taxonomic information using taxonkit (v0.20.0). We then extracted contigs whose assigned domain was the “unclassified viruses domain” (a higher-rank category encompassing all viruses). We further performed BLASTn (2.15.0+) searches against the latest NCBI core nt database available at the time of analysis (September or November 2025, or February 2026) and excluded contigs whose top BLASTn hits were assigned to non-viral sequences. This step was intended to remove sequences that may not be detected by protein-based searches but are detected by nucleotide-based searches (e.g., pseudogenized endogenous viral sequences embedded in host genomes). Contigs remaining after these filtering steps were defined as viral-derived contigs. The top-hit virus, hit length, sequence identity, and taxonomic information associated with these contigs were based on the BLASTx results against the NCBI nr database.

To focus on viruses infecting eukaryotes, we excluded bacteriophage-related taxa (*Caudoviricetes*, *Inoviridae*, *Lavidaviridae*, *Microviridae*, and virus species whose names contain “phage”) from the viral-derived contigs. In addition, to maintain consistency with ISG-VIP predictions, we excluded *Retroviridae* (which contains many endogenous retroviruses) and viral families that do not induce ISGs (*Orthoherpesviridae*, *Polyomaviridae*, *Papillomaviridae*, *Circoviridae*, *Togaviridae*, *Flaviviridae*, *Arenaviridae*, *Peribunyaviridae*, *Phenuiviridae*, *Bornaviridae*, *Hepadnaviridae*, *Poxviridae*, *Adintoviridae*, and *Adenoviridae*). Details of viral-derived contigs identified in this study are provided in **Data set 12.**

### Detection of infection events and candidate infection events by novel viruses

We partitioned the viral-derived contigs identified into “viral infection events,” defined as unique combinations of RNA-Seq data (SRA Run ID) and assigned viral genus. For each infection event, we obtained (i) the cumulative amino-acid length of BLASTx hit regions, (ii) the mean sequence identity, and (iii) the modal viral species, all based on BLASTx against the NCBI nr database. Finally, infection events with cumulative hit length > 200 amino acids and mean top hit sequence identity < 95% were defined as candidate infection events by novel viruses.

### Genome alignment by TBLASTx

To evaluate which genomic regions of reference viruses were covered by the detected viral-derived contigs, we mapped contigs to known reference viral genomes using TBLASTx.

To select reference viral genomes for individual viral infection events, we first extracted the top-hit viral proteins from the BLASTx-based virus search above. Using these proteins as queries, we performed TBLASTn searches via the NCBI BLAST web interface. Among the resulting subjects, we selected nucleotide sequences that covered nearly the full length of the viral genome (or the corresponding segment for segmented viruses) as the reference viral sequences. Gene annotations for the selected reference sequences were also retrieved from NCBI.

Contigs assembled from RNA-seq data corresponding to each infection event were then mapped to the appropriate reference genomes using the tblastn command in BLAST+ (v2.17.0). We set word size to 3 and the e-value threshold to 1.0e-4, and used default settings for all other parameters.

### Phylogenetic analysis based on high-quality contigs assembled by Trinity

Details of the phylogenetic trees estimated in this study (target viral taxonomy and protein, number of sequences, thresholds for quality control, etc.) are summarized in **Table S6**. To build a reference MSA, we retrieved all amino-acid sequences for the target viral taxon and protein from NCBI Virus (https://www.ncbi.nlm.nih.gov/labs/virus/vssi/#/). We excluded sequences with >10% undetermined amino acids and sequences shorter than the data set-specific length threshold. Sequences lacking host information were also removed. We next removed redundant sequences by clustering with CD-HIT (v4.8.1)^53^ using a data set-specific threshold based on pairwise p-distance. We then constructed a reference MSA using MAFFT (v7.490; --auto)^54^. In parallel, the longest ORFs (flanked by stop codons) were extracted from Trinity-assembled virus-derived high-quality contigs and translated into amino acid sequences. Subsequently, we added the contig-derived amino acid sequences to the reference MSA using the --addfragments option. Using TrimAl (v1.5.rev0)^55^, we removed alignment sites with gaps in >50% of sequences. We further manually trimmed terminal regions absent from the target contig-derived amino acid sequences and removed clearly inappropriate sequences (e.g., sequences for which most alignment sites were gaps) to obtain the final MSA. Maximum likelihood phylogenetic trees were then inferred using iqtree3 (v3.0.1)^56^. Model selection was performed using ModelFinder^57^, and site-rate heterogeneity was modeled using the FreeRate model. Branch support was assessed with 1,000 ultrafast bootstrap replicates. Details of the sequence data used in the phylogenetic analyses are listed in **Data set 13**.

### Phylogenetic analysis based on Logan contigs

This analysis was performed to construct the pan-parvoviral phylogenetic tree. Because Logan contigs tended to be short and fragmented, we generated contig assemblies by concatenating contigs guided by the reference MSA and used them for phylogenetic inference.

For each infection event we extracted contigs that hit the target protein by BLASTx using amino acid sequences in the reference MSA as the database. Next, the longest ORFs (flanked by stop codons) were extracted from the virus-derived contigs and translated into amino acid sequences. Subsequently, we mapped them onto the reference MSA using MAFFT --addfragments. Contig-derived amino acid sequences were sorted based on their mapped regions on the reference MSA, and overlaps between adjacent sequences were evaluated. If the overlap exceeded 5% of the alignment length, the shorter contig-derived amino acid sequence was removed. The remaining contig-derived amino acid sequences were concatenated in the sorted order to generate an assembly sequence, which was used as a substitute for the high-quality contig-derived amino acid sequences in phylogenetic analysis.

We applied additional filtering to the trimmed MSA: we removed sequences with >30% gaps among known reference sequences and sequences with >60% gaps among the newly assembled sequences. To exclude highly divergent outliers lacking similarity to other sequences in the data set, we computed pairwise p-distances within the MSA and removed sequences whose minimum (nearest-neighbor) p-distance to any other sequence exceeded 0.85 prior to phylogenetic inference.

### Ancestral host inference

We inferred ancestral hosts based on ancestral discrete-trait reconstruction. We used the maximum likelihood phylogenetic tree estimated from Trinity-based high-quality contigs. Using make.simmap in phytools (v2.5.2)^58^, we performed Markov-model-based ancestral discrete-trait reconstruction with the all rate different (ARD) transition model and a root prior estimated from the data. Host family was used as the discrete trait. We performed 1,000 stochastic mapping replicates and obtained marginal posterior probabilities of each trait at ancestral nodes.

### Quantification of reads derived from a novel *Betaarterivirus* in reed vole RNA-seq data

We first constructed a custom reference genome database that included the reed vole reference genome and the longest contig of the newly identified *Betaarterivirus*. We also generated a corresponding custom gene annotation by incorporating the novel *Betaarterivirus*, treating the full-length viral genome sequence as a single exon. Using a standard RNA-seq pipeline with STAR and featureCounts, we quantified RPM for both host genes and viral genes. The reference genomes and gene annotations used are listed in **Table S1**, and the RNA-seq data sets analyzed are provided in **Table S3**.

### Differential expression and GSEA

We performed differential expression analysis between infected and uninfected samples using DESeq2. Using NCBI gene ortholog information, we mapped genes of the target animal species to human orthologs. Based on human gene symbols and Wald statistics from DESeq2, we performed pre-ranked GSEA using gseapy (v1.1.11)^59^. We used MSigDB Hallmark 2020^60^ as the gene set collection.

## Code availability

The computational code used in this study and the ISG Profiler/VIP package are available at the GitHub repository (https://github.com/TheSatoLab/ISG-Profiler_VIP).

## Data availability

The following datasets are deposited in the Zenodo repository (https://zenodo.org/records/19045649).

## Acknowledgments

The super-computing resource was provided by Human Genome Center (the Univ. of Tokyo). ISG Profiler/VIP was packaged by Genta Kuroki at Youworks Corporation.

This study was supported in part by JST PRESTO (JPMJPR22R1, to Jumpei Ito); JSPS KAKENHI Grant-in-Aid for Scientific Research B (JP25K00116, to Jumpei Ito); AMED SCARDA Japan Initiative for World-leading Vaccine Research and Development Centers “UTOPIA” (JP223fa627001, to Jumpei Ito, Kei Sato, Spyros Lytras, Luca Nishimura); SHIONOGI Infectious Disease Research Promotion Foundation (to Jumpei Ito); JSPS KAKENHI Fund for the Promotion of Joint International Research (International Leading Research) (JP23K20041, to Kei Sato); AMED ASPIRE Program (25jf0126002, to Kei Sato); AMED SCARDA Program on R&D of new generation vaccine including new modality application (253fa727002, to Kei Sato); AMED Research Program on Emerging and Re-emerging Infectious Diseases (24fk0108907, 25fk0108690, to Kei Sato); AMED Japan Program for Infectious Diseases Research and Infrastructure (Collaborative Research via Overseas Research Centers) (25wm0225041, to Kei Sato); JSPS KAKENHI Grant-in-Aid for Scientific Research A (JP24H00607, to Kei Sato); the Platform Project for Supporting Drug Discovery and Life Science Research (Basis for Supporting Innovative Drug Discovery and Life Science Research (BINDS)) from AMED (JP24ama121012, supporting numbers S02820001 and S02820002, to Kei Sato); JSPS KAKENHI Research Fellowships for Young Scientists, PD (24KJ0068; to Luca Nishimura); JSPS KAKENHI Grant-in-Aid for Early-Career Scientists (25K18449; to Luca Nishimura); National Health and Medical Research Council (Australia) Investigator grant (GNT2017197 to Edward C. Holmes).

## Author contributions

Project design, management, and supervision: Jumpei Ito

Development and validation of ISG Profiler: Luca Nishimura

Preparation of ISG expression and viral detection profiles from large-scale public RNA-seq data: Mai Suganami and Luca Nishimura

Comparative analysis of ISG expression: Luca Nishimura

Development and validation of ISG-VIP: Hiroaki Unno

Virus discovery and characterization: Kyoko Kurihara and Jumpei Ito

Evolutionary analyses: Jumpei Ito

Computational cost evaluation: Mai Suganami

Data set provision: Junna Kawasaki

ISG-Profiler/ISG-VIP package development: Jumpei Ito

Manuscript drafting: Jumpei Ito

Manuscript edit: Jumpei Ito, Edward C Holmes, Spyros Lytras, Luca Nishimura, Junna Kawasaki, and Kyoko Kurihara

Figure preparation: Jumpei Ito and Luca Nishimura

Scientific illustrations: Jumpei Ito and Kaho Okumura

Logo design: Luca Nishimura and Kaho Okumura

Bioinformatics analysis support: Mai Suganami, Jumpei Ito, Kaho Okumura, Junna Kawasaki, Spyros Lytras, Edward C Holmes, and Kei Sato

All authors reviewed and approved the final manuscript.

## Supplemental Table

Table S1: Reference genome and gene annotation information used for STAR mapping.

Table S2: List of ISGs and ICGs used in ISG Profiler.

Table S3: RNA-seq data sets used for detailed analyses.

Table S4: Information on viral infection events detected by BLASTx. Table S5: Information on reassembled high-quality contigs.

Table S6: Information on parameters used in phylogenetic analyses.

## Data sets

Data set 1: Metadata for the Shaw et al. RNA-seq data set.

Data set 2: Metadata for the He et al. and Zhao et al. RNA-seq data sets.

Data set 3: Metadata for the 170k RNA-seq data set.

Data set 4: Metadata for the 40k RNA-seq data set.

Data set 5: ISG Profiler output for the Shaw et al. RNA-seq data set.

Data set 6: ISG Profiler output for the He et al. and Zhao et al. RNA-seq data sets.

Data set 7: ISG Profiler output for the 170k RNA-seq data set.

Data set 8: ISG Profiler output for the 40k RNA-seq data set.

Data set 9: Metadata for sequences in the ortholog database.

Data set 10: geNomad results for the 170k RNA-seq data set.

Data set 11: geNomad results for the 40k RNA-seq data set.

Data set 12: Viral-derived contigs identified by BLASTx.

Data set 13: Metadata for viral sequences used in the phylogenetic analyses.

## Supplemental figure legends

**Fig. S1. Development of ISG Profiler**

A) ISG Profiler computational time. Twenty paired-end 150-bp RNA-seq data set derived from diverse animal species were analyzed on a computational unit equipped with an AMD EPYC 9654 (2.4 GHz) processor running Red Hat Enterprise Linux 8 HPC (x86_64), using 8 CPU cores and 16 GB of RAM.

B) Increase in ISG scores upon IFN treatment in the RNA-seq data set from Shaw et al.^13^. Boxplots show quartiles and Tukey’s fences. Asterisks indicate *P* < 0.05 (two-sided Welch’s *t*-test).

C) Changes in ISG scores associated with natural viral infection. RNA-seq data from lung samples of diverse animal species with previously reported viral infections (He *et al.* and Zhao *et al.*^20,21^) were analyzed. Heatmap colors indicate differences in mean ISG scores between virus-positive and virus-negative samples for each unique combination of animal species and viral family. Dot size reflects the number of samples.

D) Ablation analysis assessing the robustness of ISG Profiler to limited genome availability focusing on chickens. Top: ISG scores for IFN-treated and untreated samples calculated using the original and reduced databases; dashed lines indicate medians obtained with the original database. Middle: Differences in mean ISG scores between IFN-treated and untreated samples. Bottom: Composition of each ortholog database, showing the taxonomic levels of reference sequences included.

E) Increase in ISG scores following experimental viral infection in animal species lacking species-specific reference sequences. For *Chlorocebus aethiops* and *Sigmodon* species, reference sequences were available only at the genus and family levels, respectively. Asterisks indicate *P* < 0.05 (two-sided Welch’s *t*-test). HSV, Herpes simplex virus; RSV, Respiratory syncytial virus. Sample information is listed in **Table S3**.

F) Conservation patterns of the 59 core ISGs across Aves *and* Mammalia. The conservation rate for each order is shown. Core ISGs can be categorized into three gene clusters: mammalia-specific, placentalia-specific, and genes widely conserved across these taxa.

G) Species-specific standardized counts of each ISG across 398 amniote species, based on IFN-treated human (ERR2012472; upper) and chicken (ERR2012446; lower) RNA-seq data from Shaw et al. Rows and columns represent core ISGs and animal species, respectively. Arrows indicate the column corresponding to human or chicken.

H) Comparison of expression estimates between ISG Profiler and a conventional RNA-seq pipeline. RNA-seq data from Shaw et al. were analyzed. Dots represent core ISGs. The x-axis shows the log-transformed mean RPM estimated using a conventional reference-based pipeline (STAR and featureCounts), whereas the y-axis shows the normalized counts calculated by ISG Profiler using 100 ICGs for normalization. Solid lines indicate linear regression fits. ISGs excluded from the final set of 59 core ISGs are highlighted in color.

I) Comparison of expression estimates between ISG Profiler and a conventional RNA-seq pipeline. Results obtained using a prototype version of ISG Profiler with 30 ICGs are shown. As an example, data from *Rattus norvegicus* are presented.

J) Overview of the selection scheme for 100 ICGs. Numbers on the right indicate the number of genes retained at each selection step.

K) Selection of the mapping tool used in ISG Profiler. Using five different mapping tools, differences in standardized counts between IFN-treated and untreated samples were calculated using RNA-seq data from Shaw et al. Spearman correlations between these values and log₂ fold-change estimates obtained from a conventional pipeline (STAR, featureCounts, and DESeq2) were calculated for each mapping tool. Heatmap rows (mapping tools) are sorted by the average Spearman correlation across species.

**Fig. S2. ISG profiling at the pan-SRA scale across host species and viral taxa**

A) Number of contigs assigned to each viral family. Primary assignments were based on geNomad, and virus-like contigs were further examined using BLASTx searches against the NCBI non-redundant (nr) protein database. Bar plots are stratified by top-hit sequence identity in BLASTx. Contigs without viral hits in BLASTx are labeled “No viral hit,” and contigs assigned to different viral families by geNomad and BLASTx are labeled “Unmatched virus family.”

B) Number of samples in which each viral family assigned by geNomad was detected.

C) Composition of viral species within *Coronaviridae*. Based on BLASTx results, contigs with ≥95% top-hit identity were assigned to the corresponding viral species and included in the analysis.

D) Composition of host animal species for *Betacoronavirus pandemicum*, including SARS-CoV, SARS-CoV-2, and related coronaviruses.

E) Composition of No viral hit (i.e., false-positive) contigs assigned to *Orthoherpesviridae* by geNomad. Protein categories of the top BLASTx hits are shown.

F) Distribution of basal ISG scores across animal orders. Dots represent individual animal species. Boxplots show quartiles and Tukey’s fences. Asterisks indicate statistical significance (*P* < 0.05; two-sided Wald test in a linear mixed-effects model); asterisk color indicates positive (red) or negative (blue) effects.

G) Mean ISG scores for non-infected samples and samples infected with DNA or RNA viruses across animal species. The top 10 species with the largest numbers of virus-positive samples are shown.

H) Mean ISG scores for non-infected samples and samples infected with each viral family across animal species. The top 10 animal species with the largest numbers of virus-positive samples are shown. Dot size reflects the number of samples. Asterisks indicate statistical significance (*P* < 0.05; two-sided Wald test in a linear mixed-effects model); asterisk color indicates positive (red) and negative (blue) effects. NS, not significant.

I) Comparison of virus detection by geNomad^23^ and by Kawasaki et al.^3^, in which RNA viruses were comprehensively identified from mammalian and avian RNA-seq data in the SRA using BLASTx.

J) Differences in ISG scores during *Orthoherpesviridae* infection. The left panel shows results based solely on geNomad classification, whereas the right panel shows results based on both geNomad and BLASTx classifications.

**Fig. S3. Development of the viral infection prediction model ISG-VIP**

A) F1 scores stratified by viral family (left) and host species (right). Only viral families and host species with ≥50 virus-positive samples are shown. Mean values (bars) and standard deviations (error bars) are shown.

B) SHAP (SHapley Additive exPlanations) scores for individual variables in the LightGBM component of ISG-VIP.

C) Precision, recall, and predicted positive rate of ISG-VIP in the internal validation.

D) Comparison of virus-derived contigs generated by the Logan assembly pipeline (Cuttlefish2 + Minia3)^22,28,45^, MEGAHIT^52^, and Trinity^30^. The example shown corresponds to a *Betaarterivirus* detected in reed voles. Virus-derived contigs were aligned to the near-complete reference genome most closely related to the BLASTx top-hit virus. Each row represents a contig, and color intensity indicates amino acid sequence identity. Gene annotations of the reference genome are shown above.

E) Computational cost comparison of the Logan assembly pipeline, MEGAHIT, and Trinity. Twenty paired-end 150-bp RNA-seq data set derived from diverse animal species were analyzed on a computational unit equipped with an AMD EPYC 9654 (2.4 GHz) processor running Red Hat Enterprise Linux 8 HPC (x86_64), using 8 CPU cores and 192 GB of RAM. WData are presented as mean ± standard deviation.

**Fig. S4 Alignment of identified viral contigs to the reference genome.**

Mapping of viral contigs to the reference genome by TBLASTx. High-quality contigs obtained by Trinity^30^ reassembly were mapped to the near-complete reference genome most closely related to the BLASTx top-hit virus. Rows represent contigs, and color indicates amino acid sequence identity. Gene annotations of the reference genome are shown. Viruses identified in the external validation data set are shown. Details of reassembled high-quality contigs are listed in **Table S5**.

We identified sequences of *Betaarterivirus* and *Gammaarterivirus* in multiple rodent species. *Gammacoronavirus* sequences were detected in pigeons, and *Deltacoronavirus* sequences were identified in pigs; however, the latter corresponded to the known porcine deltacoronavirus (*Coronavirus* HKU15; *Deltacoronavirus suis*). *Hepeviridae* sequences were detected in multiple rodent species as well as in zebra finch (*Taeniopygia guttata*). An orthonairovirus, *Orthonairovirus bushkeyense*, previously reported exclusively from tick vectors, was identified in an aquatic bird, the common murre (*Uria aalge*). Within *Paramyxoviridae*, a *Jeilongvirus* was identified in deer mice (*Peromyscus maniculatus*), although the corresponding sequences were highly fragmented. *Morbillivirus* was detected in dogs; however, this signal originated from experimental infection of cultured cells with canine distemper virus. Similarly, although *Orthorubulavirus* sequences were detected in multiple artiodactyl samples, these samples were derived from cultured cells and the detected viral sequences showed high similarity to human parainfluenza virus 5, suggesting unintended laboratory infections from human sources. *Protoparvovirus* sequences were detected in pigs, rodents, and marsupials. Within *Picornaviridae*, *Cardiovirus* was identified in multiple rodent species, *Enterovirus* in pigs, and *Hepatovirus* in chickens.

**Fig. S5 Detection of previously overlooked viral infections**

A) Reasons why a subset of samples in the TP fraction were not classified as virus-positive by BLASTx-based detection. These samples were not excluded because they lacked BLASTx hits to viral proteins. Rather, their top hits corresponded to non-viral sequences in subsequent BLASTx searches against the NCBI nr database and BLASTn searches against the NCBI nt database, suggesting false-positive detections by geNomad.

B) Proportion of virus-derived contigs identified by BLASTx that were also detected by geNomad. The results are stratified by contig length.

C) Detection of ISG non-inducing viral families by ISG-VIP. In addition to the detection proportion in the FP and TN fractions, enrichment score [log_2_(proportion in FP / proportion in TN)] for the FP fraction is shown.

D) Comparison of genetic diversity between *Chaphamaparvovirus* and other parvoviral genera. The scatter plot shows clade diversity (mean pairwise p-distance within each clade) versus the number of sequences after redundancy removal using 90% sequence identity clustering.

E) Proportion of viral infection events detected by geNomad. For each parvoviral genus, the geNomad-positive rate is shown for all events, events with >200 aa cumulative hit length, and candidates for novel viral infections. Unlike the other analyses, which used geNomad v1.8 and database v1.7, this analysis used the latest versions available as of March 2026: geNomad v1.11.2 and database v1.9.

F) Maximum likelihood tree of *Chaphamaparvovirus* NS1 proteins. Viral sequences that clustered within the *Chaphamaparvovirus* clade in the pan-parvoviral phylogenetic tree (Fig. 5G) were used. In the right panel, subtrees highlighting viruses detected by ISG-VIP and their related viruses are shown. Branches of the phylogenetic tree are colored according to host taxonomic order using a parsimonious assignment rule.

G) Gene expression changes associated with *Chaphamaparvovirus* infection in chicken liver samples. Expression changes of representative genes for each GSEA term shown in Fig. 5J are displayed. The upper heatmap indicates gene–term membership.

**Fig. S6 Characterization of newly identified viruses.**

A) Comparison of genetic diversity between the *Hepatovirus*-like avian clade and closely related picornavirus genera. Pairwise p-distances were calculated among sequences within each group, and their distributions are shown as density plots.

B) Gene expression changes associated with *Hepatovirus* infection in chicken liver samples. Expression changes of representative genes for each GSEA term shown in Fig. 6C are displayed.

C) Ancestral state reconstruction to infer the origins of FPV, CPV-2, and PPV-1. Results based on both full tree and a reduced tree, in which newly identified rat viruses were removed, are shown. Host families other than those of primary interest were grouped into an “Others” category after estimation. In the full phylogeny, the ancestral host of the clade of FPV, CPV-2, and PPV-1 was inferred with high likelihood to be Muridae. In contrast, this likelihood was substantially reduced in the phylogeny excluding the newly identified sequences, highlighting the importance of the identified sequences for inferring the evolutionary origin of this clade.

D) Magnified view of the phylogenetic tree shown in Fig. 6G. With the exception of PRRSV, the majority of *Betaarterivirus* sequences—including *Betaarterivirus chinrav* (Chinese “rat” arterivirus 1)—are derived from Cricetidae hosts rather than Muridae hosts.

