## Supplementary figures and images for "Host innate immune response profiling reveals hidden viral infections across diverse animal species"

### Fig. S1

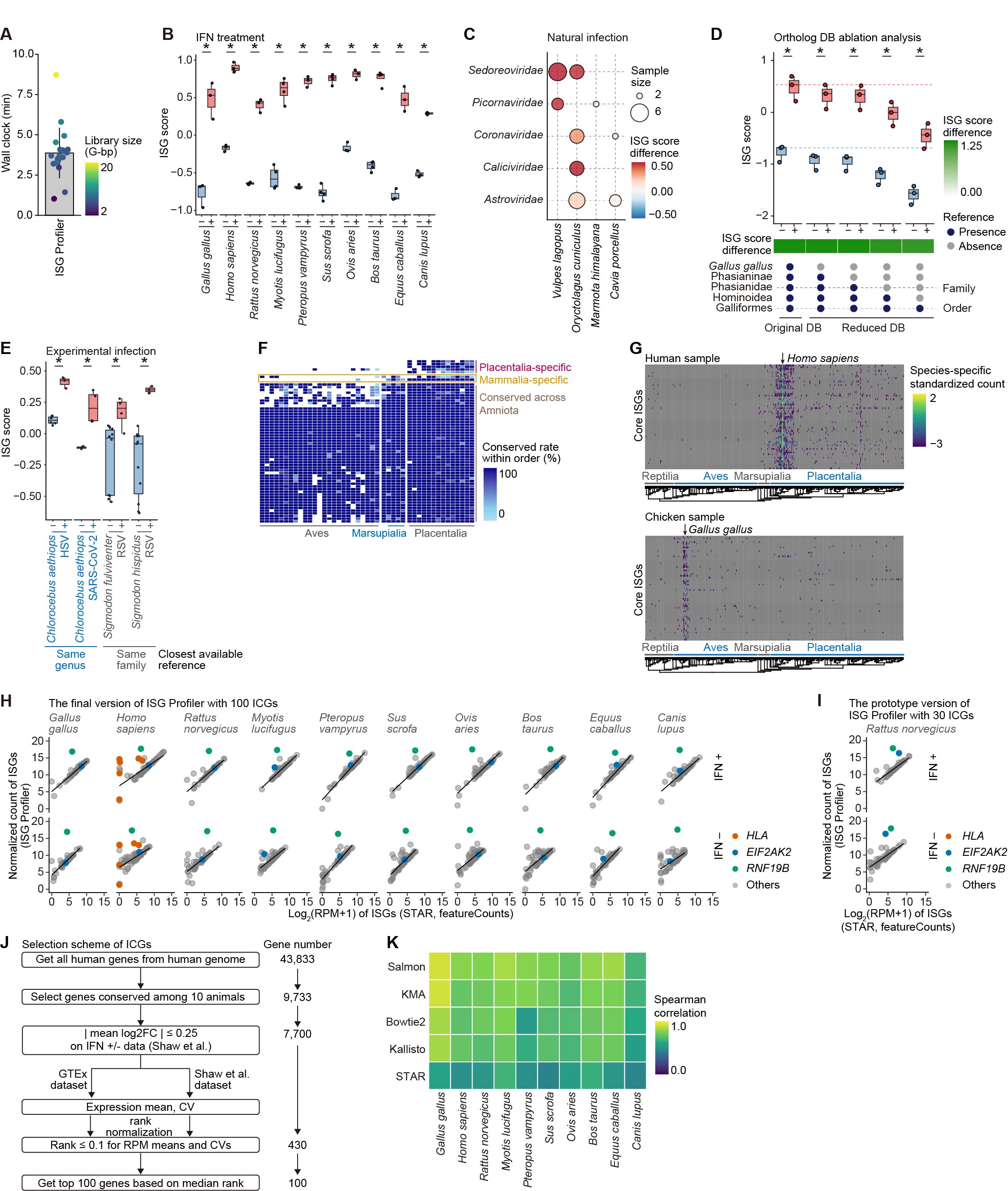

### Fig. S2

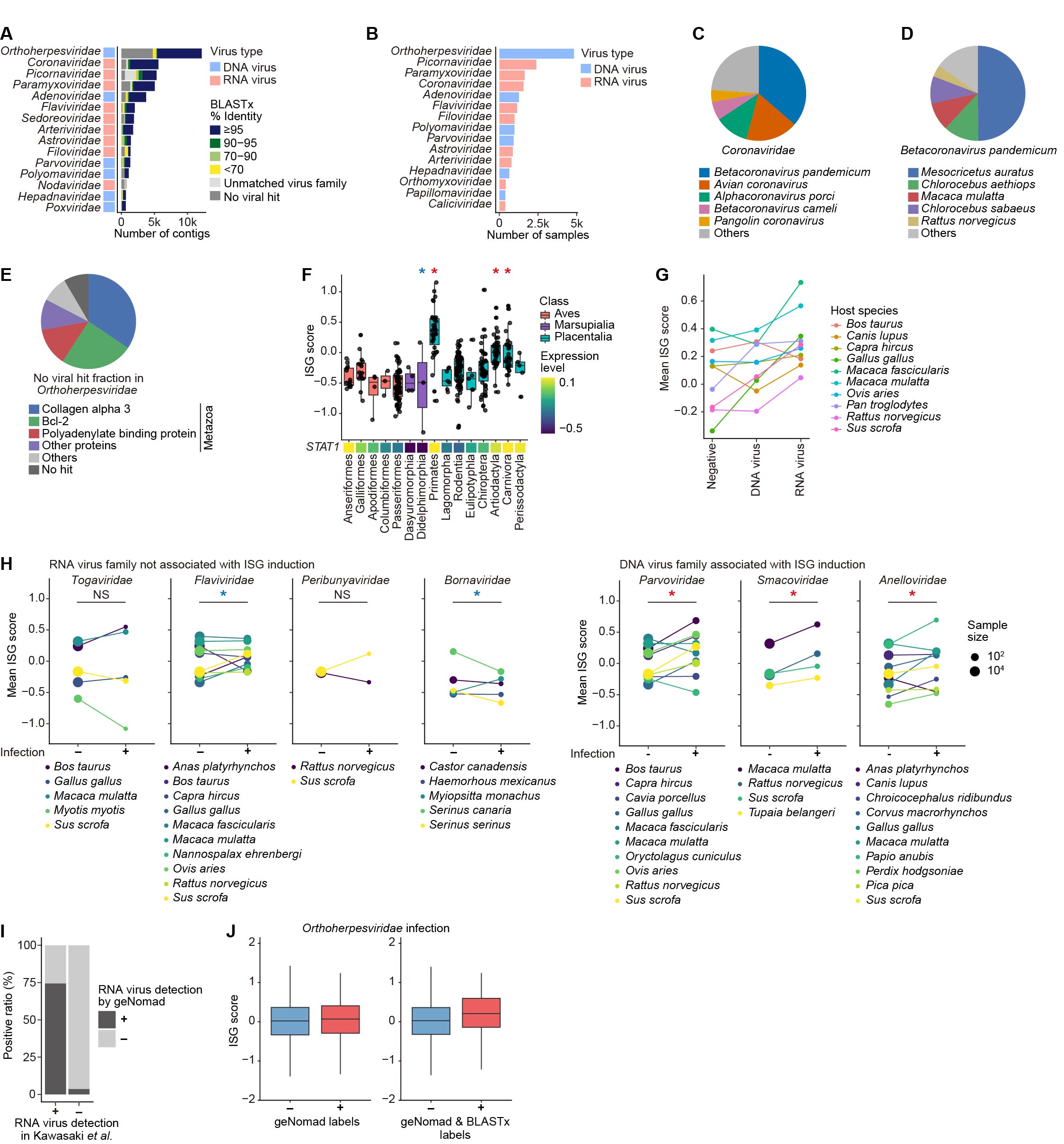

### Fig. S3

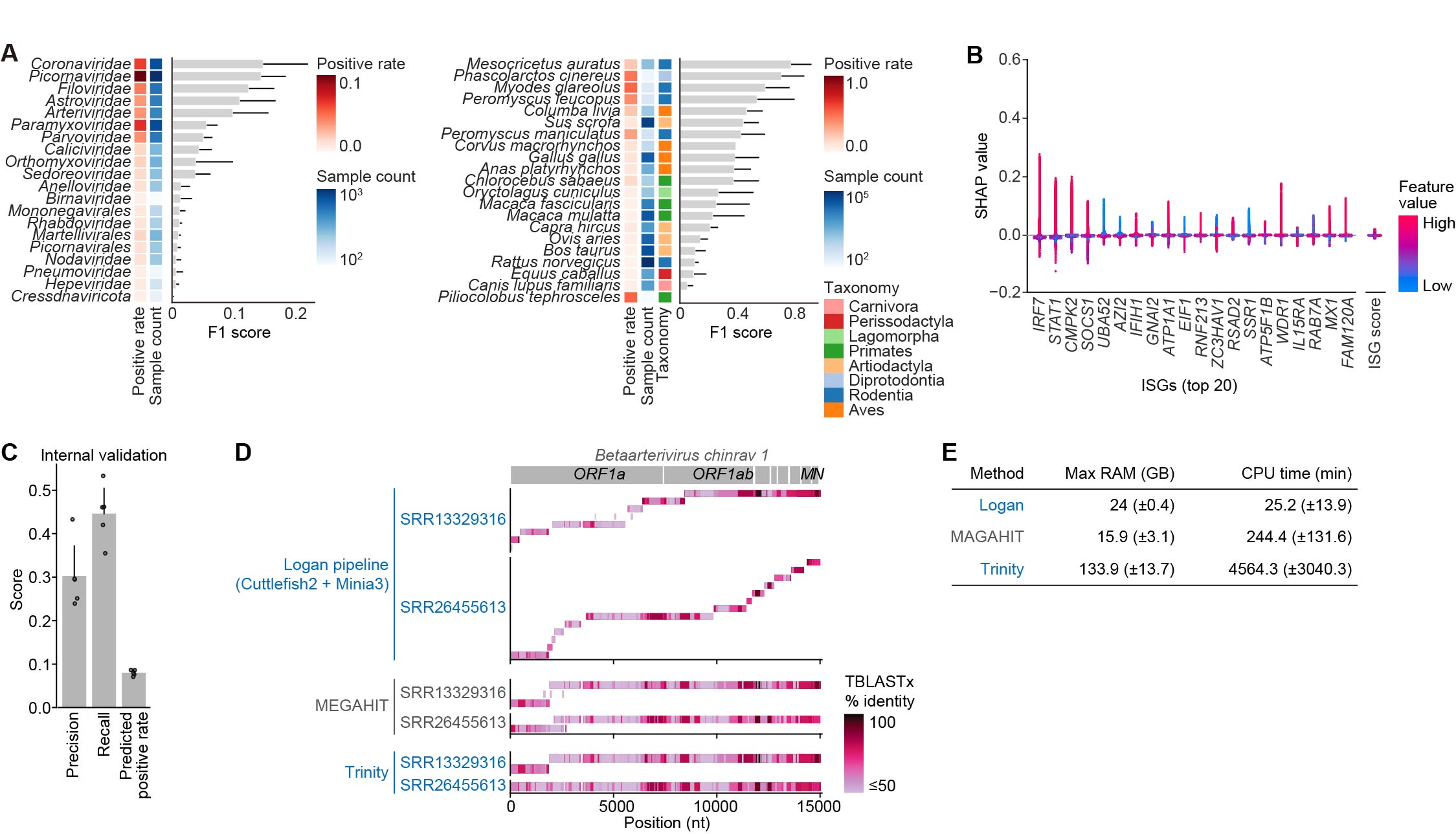

### Fig. S4

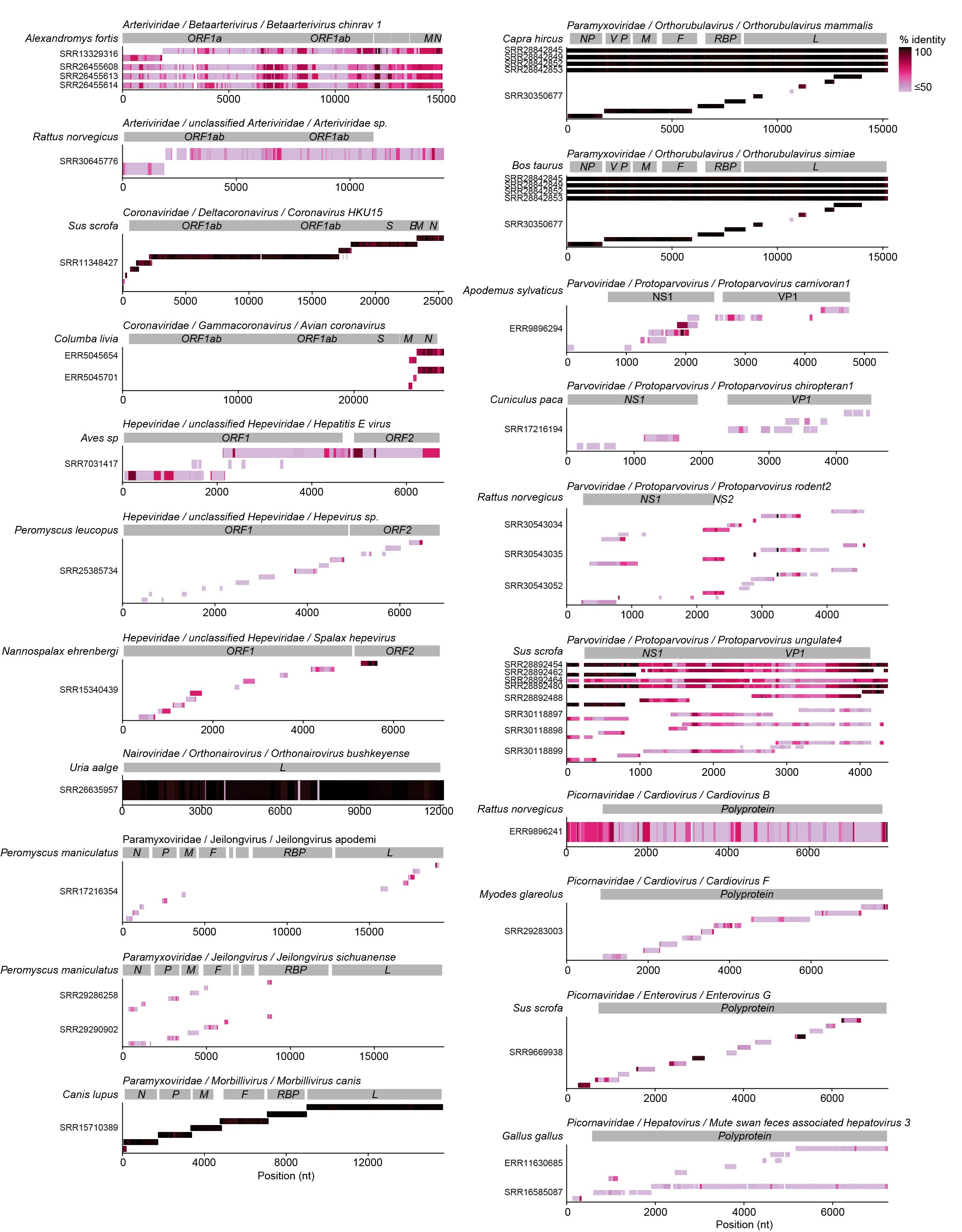

### Fig. S5

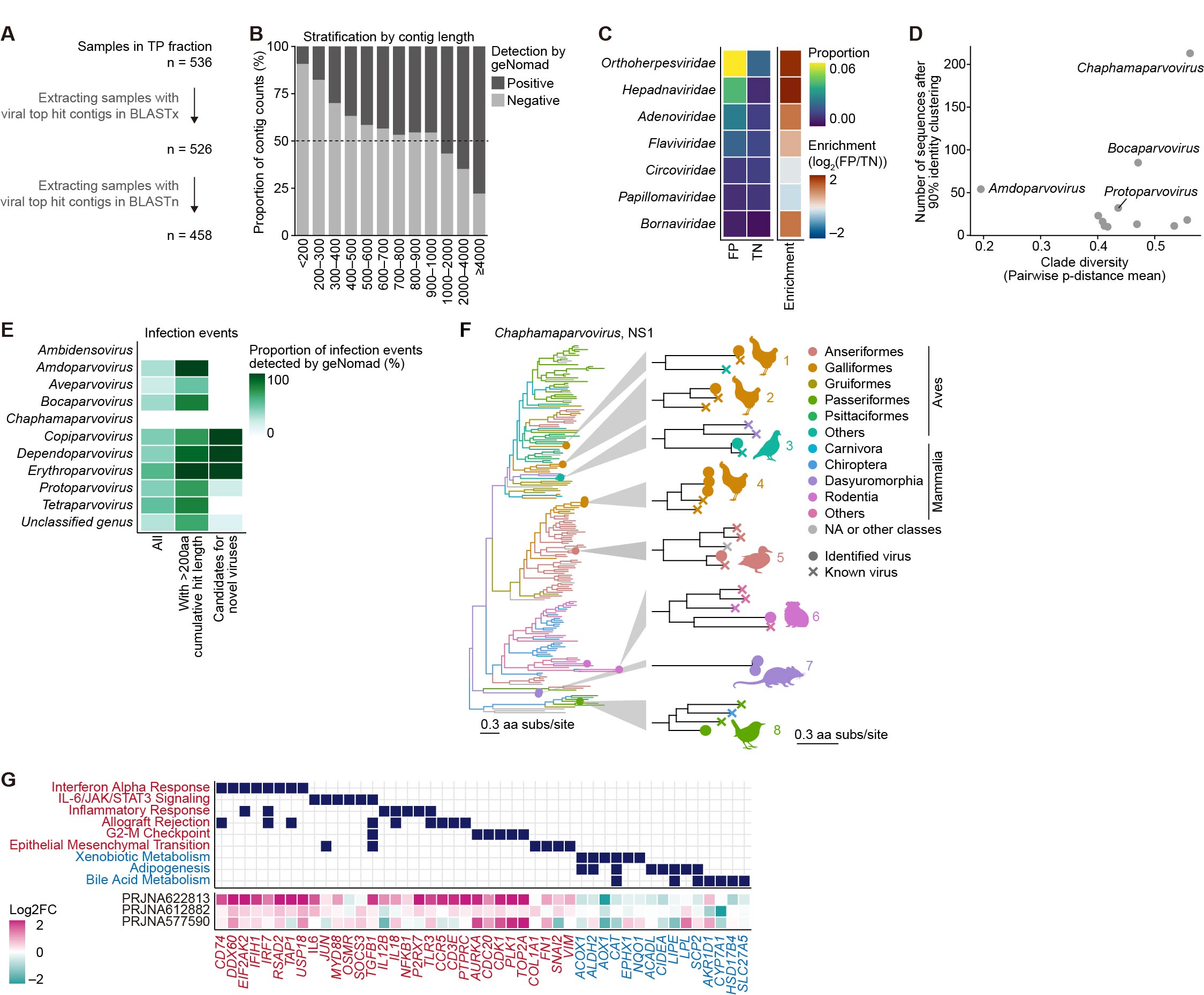

### Fig. S6

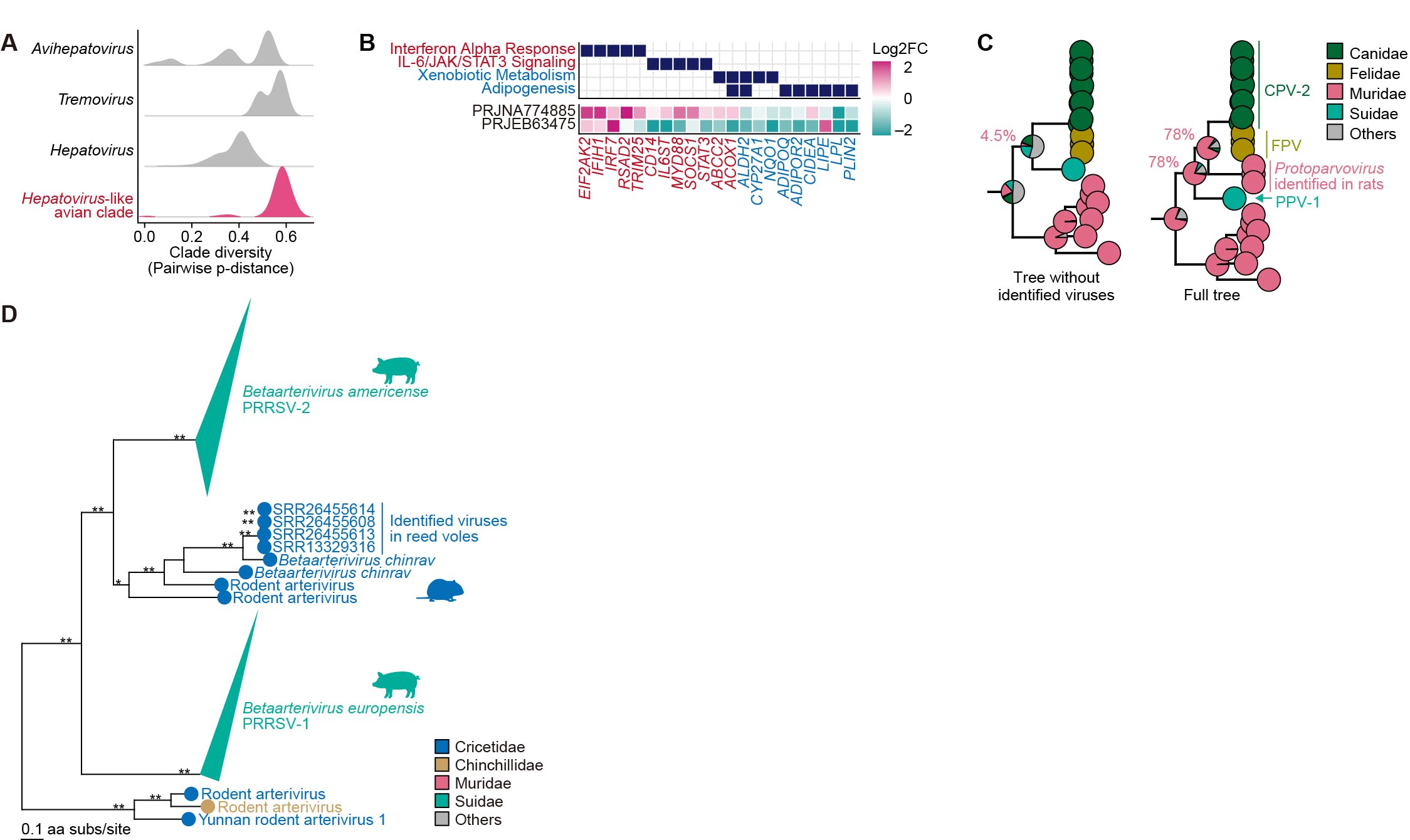
